# Cortico-cerebellar coordination facilitates neuroprosthetic control

**DOI:** 10.1101/2022.07.07.499221

**Authors:** Aamir Abbasi, Rohit Rangwani, Daniel W. Bowen, Andrew W. Fealy, Nathan P. Danielsen, Tanuj Gulati

## Abstract

Temporal coordination among neurons and development of functional neuronal assemblies is central to nervous system function and purposeful behavior. Still, there is a paucity of evidence about how functional coordination emerges in task-related neuronal assemblies in cortical and subcortical regions that are related to the control of functional output. We investigated emergent neural dynamics between primary motor cortex (*M1*) and the contralateral cerebellar cortex as rats learned a neuroprosthetic/ brain machine interface (BMI) task. BMIs offer a powerful tool to causally test how distributed neural networks achieve specific neural activation. During neuroprosthetic learning, actuator movements are causally linked to primary motor cortex (*M1*) neurons, i.e., *“direct”* neurons, that drive the decoder and whose firing is required to successfully perform the task. However, it is unknown how task-related *M1* activity interacts with cerebellar activity. We observed a striking 3–6 Hz coherence that emerged between these regions’ local–field potentials (LFPs) with neuroprosthetic learning which also modulated task-related spiking. We found a robust task-related indirect modulation in the cerebellum, and we found that this activity developed a preferential relationship with *M1* task-related direct and indirect activity but not with M1 task unrelated activity with learning. We also performed optogenetic inhibition of cerebellar activity (in the cerebellar cortex and its deep nuclei) and found that this led to performance impairments in *M1*–driven neuroprosthetic control. Together, these results demonstrate that coordinated neural dynamics emerge in cortico-cerebellar regions during neuroprosthetic learning which supports task-relevant activity in *M1* neuronal populations, and further, that cerebellar influence is necessary for *M1*–driven neuroprosthetic control.

## Introduction

To accomplish even the simplest of tasks, the nervous system coordinates activity across distant brain regions. For example, holding a bottle full of water requires several neurons to produce well-calibrated muscle forces for grasping and monitoring sensory feedback. Parallel processes in several sensorimotor regions underlie even the most trivial tasks, such as mentioned above. Dense reciprocal connectivity between these regions supports this processing which likely need to be configured rapidly and flexibly to support a repertoire of behavior and afford learning of new skills. Two motor regions, the primary motor cortex (*M1*) and the cerebellum, have dense reciprocal connections, and are known to be involved in motor learning^1, 2^. Studies have shown how learning alters local activity in the cerebellum or *M1*^3–6^, yet learning is also known to alter task-related, cross-area coordination^2, 7^. Both *M1* and the cerebellum have dense connections with other cortical and subcortical regions, and hence it is difficult to ascertain if changes in interaction are due to reciprocal connectivity between *M1* and cerebellum^8–10^, or because of their roles in coordinating a common target – the upper limb.

To isolate the effects of *M1* and cerebellum to movement control versus the direct task–related coordination, we have used a brain-machine interface (BMI) task, where select *M1* neurons (*‘direct’* neurons) modulate their activity to control an external disembodied actuator. This offers experimenters a powerful paradigm where they can dictate or set the neuron–behavior relationship. During “brain control,” direct *M1* neurons change their firing properties as the neuroprosthetic task is learned^11–18^. In addition, other neurons in the local *M1* network also become task–coupled (*i.e.*, task-related *‘indirect’* neurons)^12, 13, 15, 19–23^. It remains unknown what activity emerges in the cerebellum with *M1*–driven neuroprosthetic learning and what role it plays. Here, we hypothesized that cerebellum neurons will develop task–related firing during *M1*–driven neuroprosthetic learning. We also predicted that optogenetically inhibiting the cerebellum will impact *M1* driven neuroprosthetic control. Notably, *M1* projects to the cerebellar cortex chiefly through the cortico-ponto-cerebellar (CPC) pathway^1, 24^ and the cerebellum’s primary outflow back to the neocortex is via its deep nuclei, *i.e.*, the dentato-thalamo-cortical (DTC) pathway^25, 26^. Optogenetic manipulation studies that have targeted either the pons^24^, the deep nuclei within the cerebellum^25^, or the thalamic inputs to *M1*^26^ during rodent reaching tasks have shown that perturbations in these areas impair the reaching behavior as well as the neural dynamics in target projection area. Due to the reciprocal connectivity between *M1*–cerebellum, within the cerebellum resides the cause of *M1* activation (in its deep nuclei) as well as the consequence of *M1* activation (in the cerebellar cortex’s input layers). Hence, we performed optogenetic inhibition at both levels in the cerebellum, *i.e.*, its cortex and deep nuclei and studied the effects on *M1*–driven brain control. The chief hypothesis that we tested in this study was that cerebellum develops a task-related indirect modulation and this activation is needed for *M1*–driven BMI control.

Another focus of our investigations was on co-emergent synchronous activity across *M1* and cerebellum with neuroprosthetic learning. Recent theories have proposed that alterations in the pattern of synchronous activity across regions can serve to coordinate network activity for natural and neuroprosthetic behaviors^20, 27^. Such transient LFP activity can modulate excitability of cell groups across varying spatiotemporal scales^28, 29^. This helps achieve precise temporal control in neural networks that can enhance information transfer in specific cell populations^30^ and can influence spike-timing-dependent plasticity^31^. Such temporally coordinated activity among ensembles underlies diverse neural processes ranging from perception, decision-making, action, memory, and attention^6, 32–36^. Importantly, spiking in one region becomes coordinated with LFP in another region, indicative of synchrony^33, 37^. Despite the evidence that synchronous LFP is related to learning^6, 38^, there is a paucity of evidence that this selectively modulates task–relevant activity of neurons across brain areas. Using the neuroprosthetic task paradigm where we can control the neurons that are linked to behavioral output, we aimed to disentangle the synchronous activity of task-related *‘direct’* neurons within *M1* locally and with task–relevant *indirect* activity in the *M1* and the cerebellum (as well as task-unrelated cells of *M1*, which served as negative control) to understand how these diverse classes of cells were modulated by coordinated LFP activity in the *M1* and cerebellum.

In the BMI paradigm that we used, a small set of *M1 ‘direct’* neurons controlled a simple 1-D actuator (henceforth *M1 TR_d_* ’s)^12, 18, 21^. We recorded additional neural activity from neighboring *‘indirect’* neurons in *M1*, as well as distant cerebellar cortex. We parsed these indirect neurons as either task-related or task-unrelated neurons in the *M1* (M1 *TR_i_* ’s and M1 *TU* ’s, respectively). We looked at the relationship between these sub-classes of *M1* cells and their association to cerebellar task-related indirect activity (*i.e.*, cerebellum *TR_i_* ’s). We made the novel observation that cerebellar neurons developed strong *’task– related’* indirect modulation, and *M1* and cerebellum LFPs developed a 3–6 Hz coherence as proficient *M1*–driven neuroprosthetic control was learned. We also found that *M1 TR_d_* ’s and *TR_i_* ’s and cerebellum *TR_i_* ’s enhanced their phase-locking to this 3–6 Hz oscillation in the LFPs, but *M1 TU*s were not modulated by this 3–6 Hz LFP activity. Next, we also found that fine timescale coordination (as evaluated through canonical correlation analyses) increased between *M1* task-related neurons and cerebellar *TR_i_* ’s neurons with neuroprosthetic learning. This was not the case for *M1 TU ’*s and cerebellar *TR_i_* ’s. We also used a generalized linear model (GLM) to predict *M1 TR_d_*, *TR_i,_* and *TU* activity using cerebellar *TR_i_* activity, where we found that the cerebellar activity better predicted the *M1 TR_d_* and *TR_i_* activity, but not *M1 TU* activity. In our last set of experiments, we optogenetically inhibited the cerebellum either in the cerebellar cortex or the deep nuclei and found that inhibition of cerebellar activity at either level led to performance impairments in the neuroprosthetic task and weakening of *M1* task-related activity. Together, our results show cerebellar activation in *M1*–driven *BMI* and that the cerebellum activity develops a privileged relationship with *M1* task-related direct and indirect activity to accomplish neuroprosthetic skill learning.

## Results

We implanted microwire arrays in *M1* and silicon probes (tetrodes/ polytrodes) in the cerebellar posterior lobes (see *Methods* for details). After neural implant surgery, we trained 7 animals to exert direct neural control on the angular velocity of a mechanical actuator that can also deliver water. A linear decoder converted the firing rates of two groups of units in *M1* (randomly selected and assigned positive or negative unit weights; *M1 TR_d+_* and *TR_i–_,* respectively; *TR_d_* ’s existed only in *M1*) into the angular velocity of the actuator. We also recoded multiple other units in both *M1* and cerebellar cortex (Simplex, Crus I/II regions) that were not causally linked to the actuator movements but showed significant task–related modulation (referred as *M1 TR_i_ ’*s or cerebellum *TR_i_ ’*s). The units that did not develop task-related modulation in *M1* were classified as task–unrelated (i.e., *M1 TU*s). The number of neurons in each category per session is detailed in **Supplementary Table 1**. The *M1 TR_d_* ’s decoder gain was held constant during the session to exclusively rely on neural learning mechanisms. Each trial started with the simultaneous delivery of an auditory tone and opening of a door to allow access to the tube (**Fig. 1 a,b**). At the start of each trial, the angular position of the tube was set to resting position, *P_1_*. If the angular position of the tube was successfully controlled to go to the target position *P_2_* (see methods), a defined amount of water was delivered (*i.e., successful trial*). A trial was stopped if this was not achieved within 15s (*i.e., unsuccessful trial*). At the end of a trial, the actuator was returned to position *P_1_,* and the door was closed.

**Figure 1.**
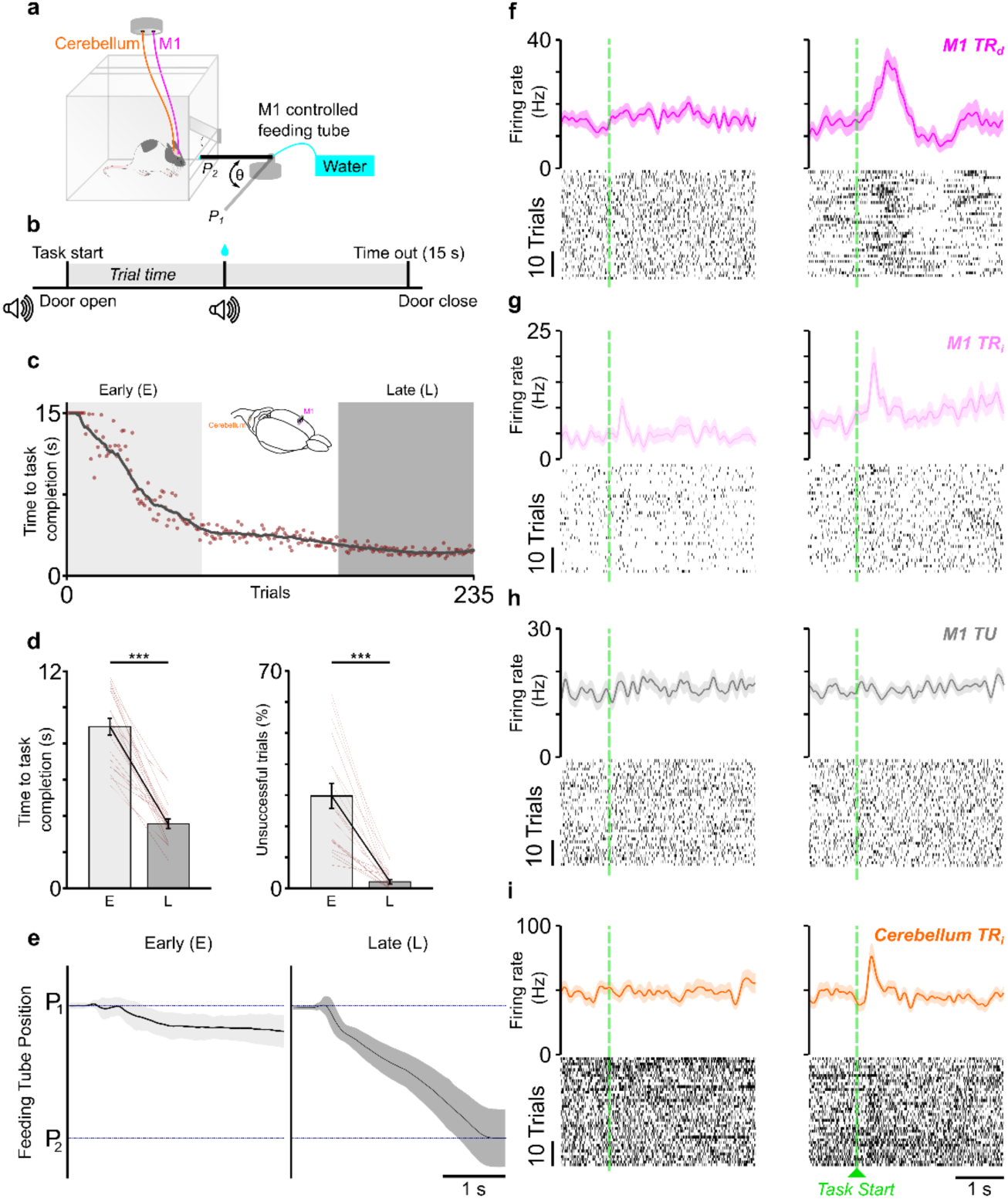
Direct and indirect modulation of *M1* and cerebellar activity with neuroprosthetic learning. ***a,*** Schematic of neuroprosthetic task box where direct neural control of a feeding tube (*θ* = angular position) was exerted. Each trial started with the tube at *P_1_*. ***b,*** Trial structure is shown depicting when audio tone cue and door movements occur. A successful trial required movement of the tube to *P_2_* within 15 seconds. ***c,*** Change in task completion time as a function of trial number from a representative session. Line shows moving average of 20 trials. Dots show individual trial task completion times. *Inset:* illustration of the recording scheme in *M1* and cerebellum from a frontal-side view. ***d,*** Change in time to task completion (*left*) and reduction in the percentage of unsuccessful trials (*right*) from *early* to *late* trials across all sessions. Bars indicate the means and error bar is s.e.m. ***e,*** Position of the feeding tube from *P_1_* to *P_2_* is shown from a single session (mean ± 1*s.d.). ***f,*** Peri-event histogram (PETH) and rasters from *early* and *late* trials from a single *M1 TR_d_* unit is shown in *left* and *right* panels, respectively. ***g,*** Same as ***f*** but for a *M1 TR_i_* unit. ***h,*** Same as ***f*** but for a *M1 TU* unit. ***i,*** Same as ***f*** but for a cerebellum *TR_i_* unit ***p<0.001.

### Direct control of BMI by M1 units

We observed that over the course of a typical 1–2 hour practice session, animals showed improvements in task performance with a significant reduction in the time to successful trial completion and decrease in the proportion of unsuccessful trials. A total of 20 sessions were analyzed, where we saw significant reductions in both metrics. Overall, we observed that rats showed improvement in task-performance with a significant reduction in the time to a successful trial completion (**Fig. 1c,d**; 8.93 ± 0.45s to 3.57 ± 0.25s, mixed-effects model: *t(38)* = –10.74, *P* = 4.5 × 10^−13^) and a decrease in the percentage of unsuccessful trials (**Fig. 1d**; 29.84 ± 4.07% to 2.19 ± 0.71%, mixed-effects model: *t(38)* = –7.03, *P* = 2.2× 10^−8^). Learning curves from all the sessions are shown in **Extended Fig. 1a**.

In a subset of sessions (*n = 13*), we tracked the position of the feeding tube, and we observed the tube’s movement from *P_1_* to *P_2_* position become direct in the *late* trials as compared to the *early* trials (**Fig. 1e**; **Extended Fig. 1b**). We also measured the speed consistency of the tube’s movement by measuring the correlation between the mean instantaneous speed (that served as template) with that of individual trials. We found that this speed consistency also increased during the *late* trials (**Extended Fig. 1c**; 0.07±0.02 to 0.23±0.03, mixed-effect models: *t(24)* = 3.86, *P* = 7 x 10^−4^). Furthermore, we also saw the angular velocity of the feeding tube was significantly higher in the *late* trials (**Extended Fig. 1d**; 5.55 ± 1.36 to 13.07 ± 2.61 pixels(px)/s, mixed-effects model: *t(24)* = 3.20, *P* = 0.0037). Using the video recording of rats during the BMI training in these same sessions, we analyzed for any correlated body movements with brain control. We looked at three movements, namely, movements of both forepaws (contralateral and ipsilateral to the implanted hemisphere) and the head. Consistent with previous reports of lack of muscle contractions during neuroprosthetic control^15^, we did not observe forepaw or head movements to be systematically correlated to the feeding tube movements, and overt movements reduced as proficient neuroprosthetic control was achieved (**Extended Fig. 2; Supplementary Video 1** and **2** from a representative *early* and *late* trial).

**Figure 2.**
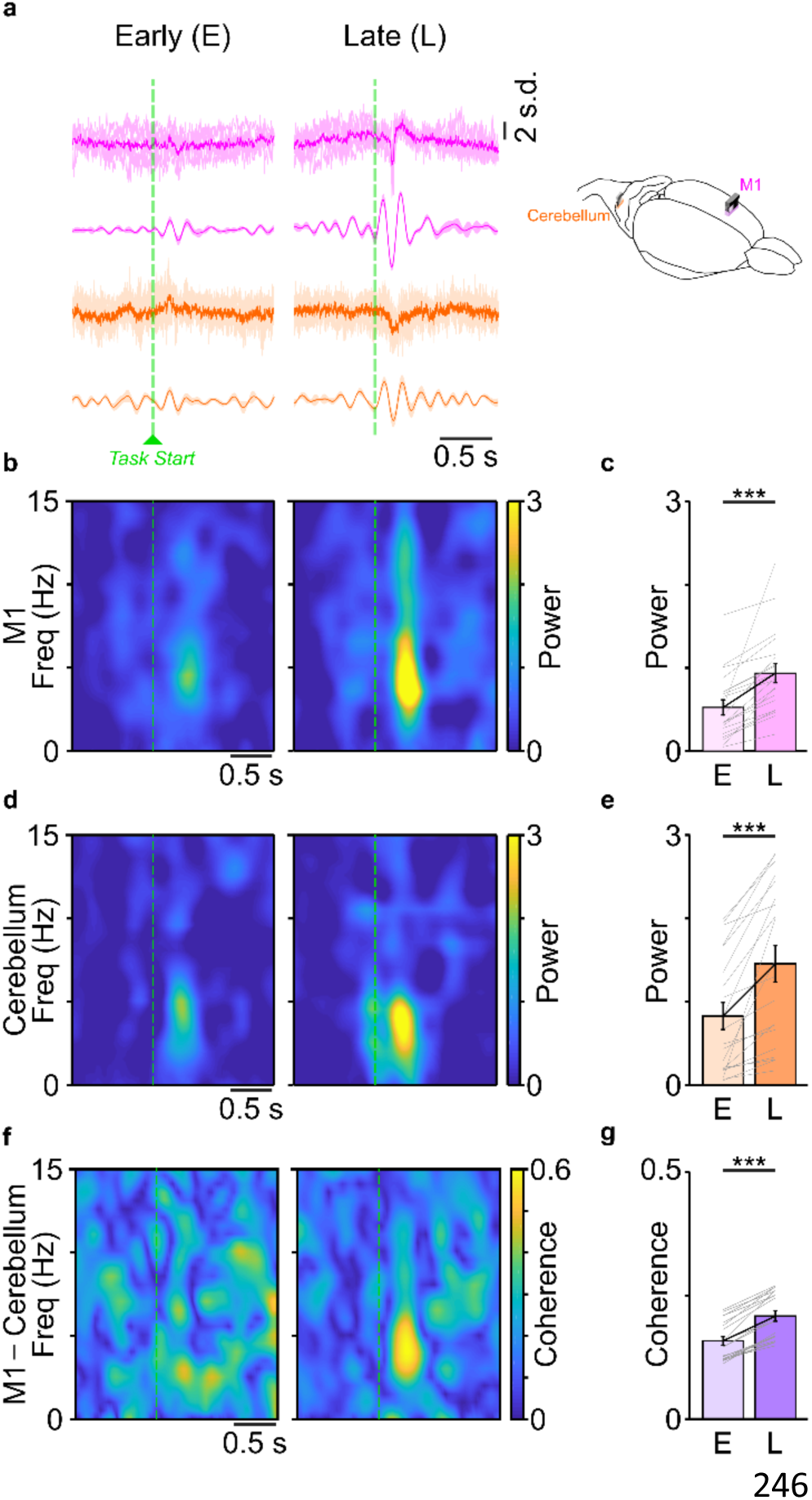
Co-ordinated neuroprosthetic task-related oscillations emerge in *M1* and cerebellar LFPs. ***a,*** Raw and filtered LFP trace from an example session showing increase in 3–6 Hz oscillations after task-start during *late* trials in both *M1* and cerebellum LFPs. Raw trace shows mean in bold overlaid on individual trial traces, filtered trace shows mean in bold and s.e.m in shaded band. *Right:* illustration of the recording scheme in *M1* (pink) and cerebellum (orange) from a frontal-side view. ***b,*** Spectrogram from a representative *M1* channel showing increase in 3–6 Hz power during *late* trials. ***c,*** Increase in 3– 6 Hz power in *M1* LFP from *early* to *late* trials across sessions. ***d,*** Same as ***b*** but from a representative cerebellum LFP channel. ***e,*** Same as ***c*** for 3–6 Hz cerebellum LFP power across sessions. ***f,*** Coherogram from a representative pair of *M1*–cerebellum LFP channel pair showing increase in 3–6 Hz coherence during *late* trials. ***g,*** Change in 3–6 Hz coherence from *early* to *late* trials across sessions. ***p<0.001.

Upon checking the modulation depth (*MD_Δ_*) of M1 decoder neurons (*TR_d_* ’s), we observed that a large proportion of *TR_d_* ’s developed robust task-related modulation (see Methods). **Figure 1f**, depicts the increase in the activity of a representative *M1 TR_d_* unit from *early* to *late* trials. Since the direct units were causally linked to the movement of the actuator, overall 90% of 89 *TR_d_* ’s showed significant *MD_Δ_* which is consistent with previous work^11^.

### Indirect modulation of neurons in M1 and cerebellum

We also analyzed indirect units in the *M1* and cerebellum and checked for their task related modulation. We further sub-classified indirect units as either task-related (*TR_i_* ’s) or task-unrelated (*TU* ’s) based on changes in *MD_Δ_* with learning. Consistent with previous reports of task-related indirect units in the *M1* likely contributing to neuroprosthetic control^11, 13, 15, 19, 21^, we found strong indirect modulation in *M1* (**Fig. 1g**). Task-unrelated units in *M1* showed no significant modulation (**Fig. 1h**). Additionally, we found robust task-related indirect modulation of units in the cerebellum (**Fig. 1i**). A majority of units in *M1* and cerebellum developed strong, indirect task–related modulation with learning (75% of 916 M1 units and 74% of 414 cerebellum units). We also looked at the change in *MD_Δ_* and found that *M1 TR_d_* underwent greater *MD_Δ_* from *early* to *late* trials as compared to *M1* and cerebellum *TR_i_* units or *M1 TU*s (*M1 TR_d_* : 134.48 ± 61.9%; *M1 TR_i_* : 61.66 ± 8.5%; *M1 TU* : –12.79 ± 4.0%; cerebellum *TR_i_* : 25.20 ± 5.5%, Kruskal-Wallis H-test, *F_3,1240_* = 246.49, *P* = 1.1 x 10^−21^, post hoc t-test showed significant difference among *M1 TR_d_*, *TR_i_* and cerebellum *TR_i_ MD_Δ_* ’s; P < 0.05).

### Emergence of coordinated task-related activity in M1 and cerebellar LFPs

Interestingly, as the rats became proficient in *M1*–driven neuroprosthetic control, we observed that a coordinated low-frequency activity (approximately 3–6 Hz, **Fig. 2a**) emerged in both *M1* and cerebellum. The task–related LFP power between 3–6 Hz increased from *early* to *late* trials in both *M1* (**Fig. 2b,c**; 0.52 ± 0.09 to 0.93 ± 0.11, mixed-effects model: *t(38)* = 3.61, *P* = 8.6 × 10^−4^) and the cerebellum (**Fig. 2d,e**; 0.83 ± 0.16 to 1.45 ±0.21, mixed-effects model: *t(38)* = 5.28, *P* = 5.4 × 10^−6^). Task-related LFP coherence (between *M1* and cerebellum LFPs) also increased in the 3−6 Hz frequency range from *early* to *late* trials (**Fig. 2f,g**; 0.15 ± 0.008 to 0.20 ± 0.01, mixed-effects model: *t(38)* = 4.72, *P* = 3.1 × 10^−5^). To ensure this was truly learning-related emergent activity and not just due to a greater proportion of unsuccessful trials early on in training, we also performed the same analysis for equal number of successful *early* and *late* trials (**Extended Fig. 3**). We found that trend of emergent 3–6 Hz task-related LFP power in both *M1* (**Extended Fig. 3a**; 0.51 ± 0.10 to 1.13 ± 0.20, mixed-effects model: *t(38)* = 3.10, *P* = 3.6 × 10^−3^) and cerebellum (**Extended Fig. 3b**; 0.81 ± 0.20 to 1.92 ± 0.33, mixed-effects model: *t(38)* = 3.66 *P* = 7.5 × 10^−4^). Task–related LFP coherence also showed an increase for an equal number of successful *early* to *late* trials (**Extended Fig. 3c**; 0.18 ± 0.011 to 0.27 ± 0.016, mixed-effects model: *t(38)* = 5.52, *P* = 2.5 × 10^−6^). This emergence of cross-region, low-frequency activity is consistent with other observations of emergence of low-frequency activity during motor skill learning across reciprocally connected neural networks^7, 20^.

### Increase in locking between spiking and LFP after neuroprosthetic learning

We next investigated the relationship between spiking activity and the low frequency LFP oscillations that we observed during robust task engagement. Spike-triggered averaging (STA) can provide an intuitive insight into how spiking is modulated by LFPs^11, 20^. We performed STA of the LFP in *early* and *late* learning, time-locked to spikes occurring either within the same region (i.e., *M1 TR_d_* / *TR_i_* / *TU* spikes to *M1* LFP and cerebellum *TR_i_* spikes to cerebellum LFP) or in the cross-region (i.e., *M1 TR_d_* / *TR_i_* / *TU* spikes to cerebellum LFP and cerebellum *TR_i_* spikes to *M1* LFP; **Fig. 3**). If spiking activity was independent of low–frequency LFP activity, then fluctuations would yield a flat average LFP. We observed an increase in the amplitude of mean LFP oscillations in both regions around the spiking of *M1 TR_d_*, *TR_i_* and cerebellum *TR_i_* units (**Fig. 3a,b,d**). This increase was not present for *M1 TU* units (**Fig. 3c**). Furthermore, we found that when we did the STA analysis during inter-trial interval (non-task period), we did not observe an increase in amplitude of mean LFP oscillations in both regions around the spiking of any class of units.

**Figure 3.**
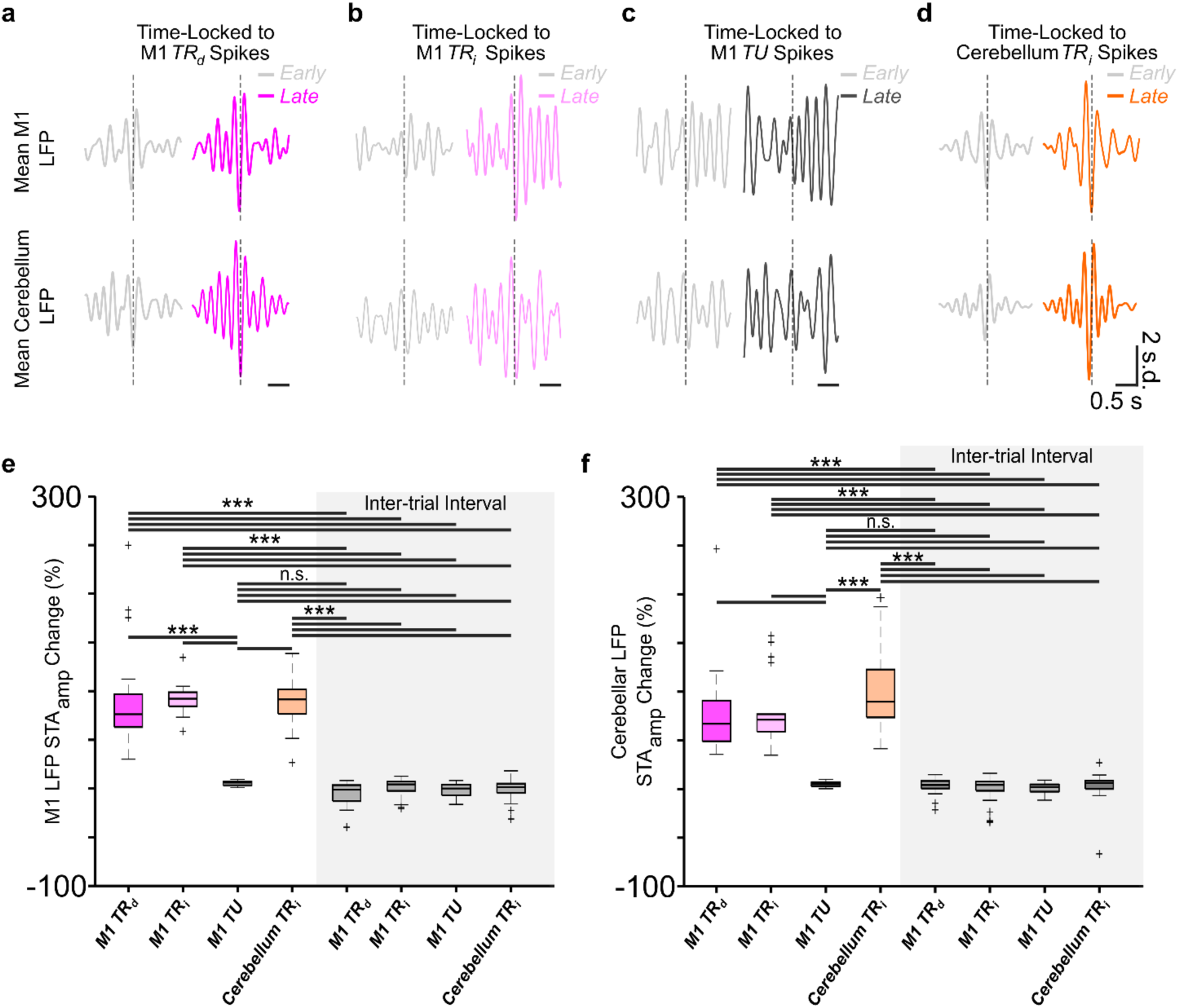
*M1* and cerebellum spike-LFP locking increases with learning. ***a,*** The mean *M1* LFP (top row) or cerebellum LFP (bottom row) time locked to occurrences of spikes from *M1 TR_d_* during task period from a representative session. ***b,*** Same as ***a*** for a *M1 TR_i_* unit. ***c,*** Same as ***a*** for a *M1 TU* unit. ***d,*** Same as ***a*** for a cerebellum *TR_i_* unit. ***e,*** Box-plot of percentage change in STA amplitude for *M1* LFP in each of the categories of units (*lower* and *upper* box boundaries are 25th and 75th percentiles, respectively, line inside the box is the median, *lower* and *upper* error lines are 10th and 90th percentiles, respectively, ‘+’ indicates outliers outside these bounds). ***f,*** Same as ***e*** for changes in STA amplitude with cerebellum LFP. ***p<0.001; n.s.: non-significant p>0.05.

*M1 TR_d_*, *TR_i_* and *TU* units, and cerebellum *TR_i_* units experienced (mean ± s.e.m.) 91.67 ± 12.39%, 91.87 ± 3.49%, 5.75 ± 0.63% and 87.31 ± 5.77% changes in STA amplitude with *M1* LFP during task periods, respectively. During inter-trial interval, these units experience −5.46 ± 2.85%, 1.08 ± 2.25%, −1.53 ± 1.57% and –1.46 ± 2.55% changes, respectively (**Fig. 3e**; Kruskal-Wallis H-test, *F_5,114_* = 90.88, *P* = 4.3 x 10^−1^^8^, post hoc *t-test* showed significant difference for *M1 TR_d_*, *TR_i_*, *TU* and cerebellum *TR_i_* during task relevant period and inter-trial period; *P* < 0.001). When STA was performed with cerebellum LFP during the task period, *M1 TR_d,_ TR_i,_* and *TU* units, and cerebellum *TR_i_* units experienced 76.25 ± 10.61%, 80.16 ± 7.84%, 4.86 ± 0.65% and 119.82 ± 25.27% changes in STA amplitude, respectively. Within area increase in STA-LFP were slightly greater than the across-area STA-LFP during task-periods. Similar to *M1* LFP, these units experienced significantly smaller change of 2.47 ± 1.99%, −0.85 ± 3.24%, 0.52 ± 1.23% and 1.45 ± 4.07%, respectively, when STA was performed with cerebellum LFP during inter-trial interval (**Fig. 3f**; Kruskal-Wallis H-test, *F_5,114_* = 91.24, *P* = 3.6 x 10^−1^^8^, post hoc *t-test* showed significant difference for *M1 TR_d_*, *TR_i_*, *TU* and cerebellum *TR_i_* during task-relevant period and inter-trial periods; *P* < 0.001).

We repeated the STA analysis for equal number of successful trials from *early* to *late* learning (**Extended Fig. 4**). *M1 TR_d_*, *TR_i,_* and *TU* units, and cerebellum *TR_i_* units experienced (mean ± s.e.m.): 95.48 ± 29.36%, 61.48 ± 9.62%, 4.64 ± 1.04% and 83.76 ± 26.31% increases in STA amplitude with *M1* LFP during task periods, respectively. During inter-trial interval (of this subset of trials), these units experienced −5.46 ± 2.85%, 1.08 ± 2.25% −1.53 ± 1.57% and –1.46 ± 2.55% change, respectively (**Extended Fig. 4a**; Kruskal-Wallis H-test, *F_5,114_* = 82.15, *P* = 2.9 x 10^−1^^6^, post hoc *t-test* showed significant difference for *M1 TR_d_*, *TR_i_*, *TU* and cerebellum *TR_i_* during task–relevant period and inter–trial period; *P* < 0.001). When STA was performed with cerebellum LFP, *M1 TR_d,_ TR_i,_* and *TU* units, and cerebellum *TR_i_* units experienced 71.90 ± 11.75%, 60 ± 11.06%, 4.98 ± 0.82% and 76.50 ± 13.31% increases in STA amplitude (during the task period), respectively. Similar to *M1* LFP, these units experienced significantly less change of 2.47 ± 1.99%, −0.85 ± 3.24%, 0.52 ± 1.23% and 1.45 ± 4.07%, respectively, when STA was performed with cerebellum LFP during inter-trial interval (**Extended Fig. 4b**; Kruskal-Wallis H-test, *F_5,114_* = 68.70, P = 1.9 x 10^−1^^3^, post hoc *t-test* showed significant difference for *M1 TR_d_*, *TR_i_*, *TU* and cerebellum *TR_i_* during task-relevant period and inter-trial periods; *P* < 0.001). With this analysis we also observed that average increase in within-area STA amplitude was greater than across-area STA amplitude.

**Figure 4.**
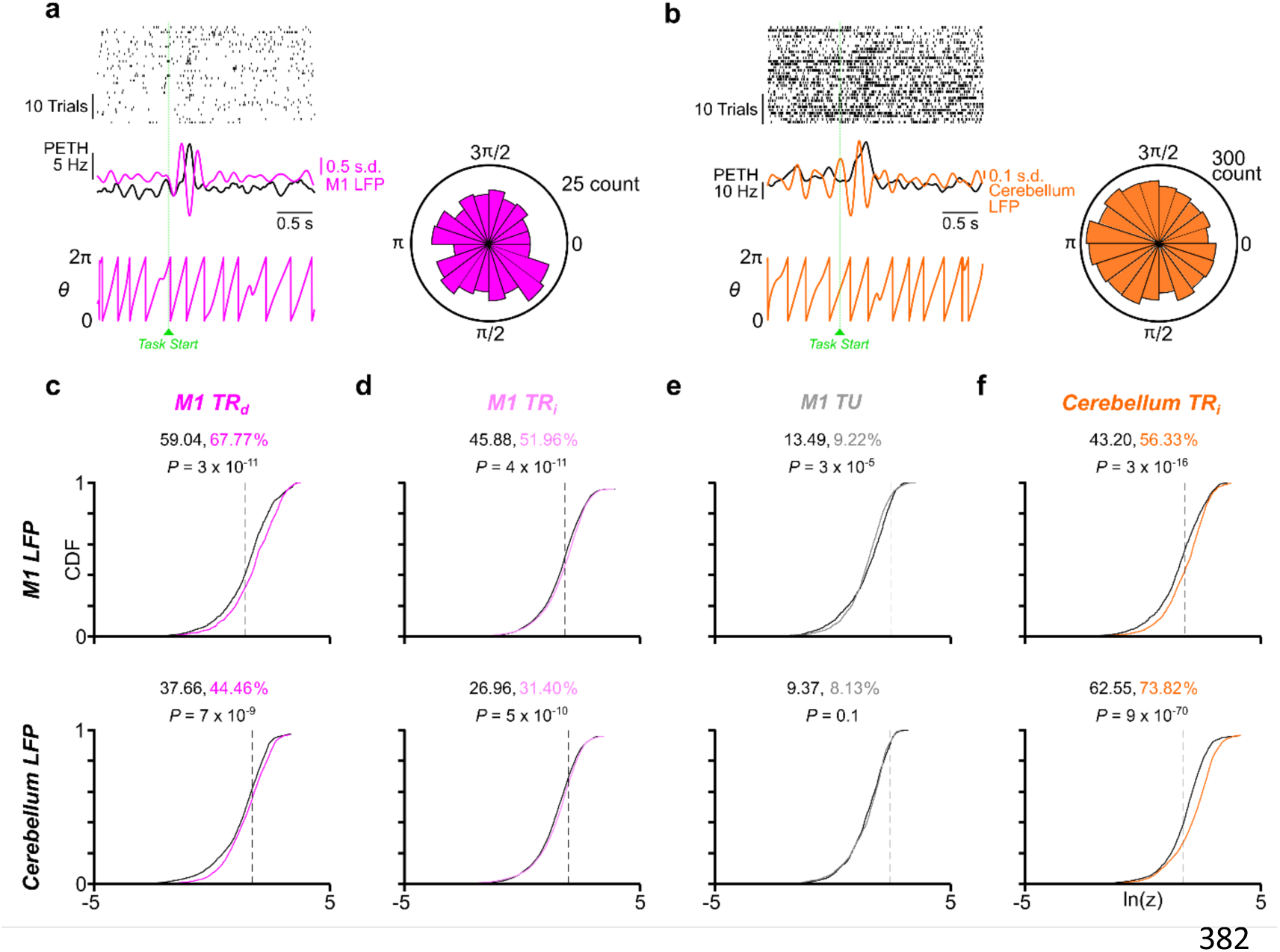
Coordinated spiking activity emerges across *M1* and cerebellum during neuroprosthetic learning. ***a,*** Spikes locking to *M1* LFP phase. *Top*: raster plot of an example *M1 TR_d_* unit. *Middle*: 3–6 Hz filtered *M1* LFP overlaid on the PETH from the example unit. *Bottom*: the extracted phase from the filtered LFP. *Right*: polar histogram from the spikes of a *M1 TR_d_* unit that showed a preferred phase to *M1* LFP. ***b,*** Same as ***a*** for a cerebellum *TR_i_* unit with a preferred phase to cerebellum LFP. ***c,*** Cumulative density functions (CDFs) of the z–statistic for every LFP–*M1 TR_d_* unit pair within and across each region. The vertical dotted lines indicate the significance threshold (*P* = 0.05) of the *z-statistic*. The percentage of the pairs with significant *P*-values is displayed. Black line indicates *early* trials, and dark pink line indicates *late* trials. The *P*-values derived using a Kolmogorov–Smirnov test. ***d,*** Same as ***c*** for *M1 TR_i_* units (light pink line indicates *late* trials). ***e,*** Same as ***c*** for *M1 TU* units (gray line indicates *late* trials). ***f,*** Same as ***c*** for cerebellum *TR_i_* units (orange line indicates *late* trials).

### Emergence of coordinated spiking–LFP activity across M1 and cerebellum

Next, we quantified phase locking of *M1* and cerebellar spikes to 3–6 Hz LFP signals in each region by generating polar histograms of the LFP phase at which each spike occurred for a single unit and LFP channel (**Fig. 4a,b**). The non-uniformity of the distribution of phases (indicating phase-locking) was quantified using a Rayleigh test of circular nonuniformity. We compared all M1 *TR_d_*, *TR_i_*, *TU* and cerebellar *TR_i_* units spiking activity from *early* to *late* trials to an M1 or cerebellar LFP channel from *early* to *late* trials. We observed an increase in the percentage of M1 *TR_d_* units that phase locked preferentially to *M1* and cerebellum LFP signals with learning (**Fig. 4c**, the black vertical dashed lines correspond to the *P* = 0.05 significance threshold of the natural log of the *z–statistic*; *M1 TR_d_* unit–*M1* LFP pairs: 59.04% to 67.77%, *P* = 3 × 10^−1^^1^, Kolmogorov–Smirnov test; *M1 TR_d_* unit–cerebellum LFP pairs: 37.66–44.46%, *P* = 7 × 10^−9^, Kolmogorov–Smirnov test). We observed that the proportion of *M1 TR_i_* units that phase locked to both *M1* and cerebellum LFPs also increased with learning (**Fig. 4d**, *M1 TR_i_* unit–*M1* LFP pairs: 45.88% to 51.96%, *P* = 4 × 10^−1^^1^, Kolmogorov–Smirnov test; *M1 TR_i_* unit–cerebellum LFP pairs: 26.98– 31.40%, *P* = 5 × 10^−10^, Kolmogorov–Smirnov test), but this was not the case for *M1 TU* units (**Fig. 4e**, *M1 TU* unit–*M1* LFP pairs: 13.49% to 9.22%, *P* = 3 × 10^−5^, Kolmogorov–Smirnov test; *M1 TU* unit–cerebellum LFP pairs: 9.37% to 8.31%, *P* = 0.1, Kolmogorov–Smirnov test). For the cerebellar *TR_i_* units, we again observed an increase in the proportion of cells that phase-locked to *M1* and cerebellum LFPs (**Fig. 4f**; cerebellum *TR_i_* unit–*M1* LFP pairs: 43.20% to 56.33%, *P* =3 x 10^−1^^6^, Kolmogorov–Smirnov test; cerebellum *TR_i_* unit–cerebellum LFP pairs: 62.55% to 73.82%, *P* = 9 × 10^−70^, Kolmogorov–Smirnov test). Notably, the task-related units of *M1* and cerebellum showed more phase-locking to *M1* or cerebellum LFPs than the task-unrelated cells of *M1*. These results indicate that the low-frequency activity that we observed emerges across *M1* and cerebellum LFPs during neuroprosthetic learning, selectively modulated task-related direct or indirect units to a greater proportion.

### Fine timescale coordination of M1 and cerebellum activity with task learning

While our analyses so far show coordinated activity in *M1* and cerebellum with task learning, it doesn’t necessarily indicate that the neural activity in the two structures were coordinated across trials. Recent studies have explored fine-timescale coordination at the level of spiking^39, 40^. Such studies use statistical methods to measure ‘communication subspaces’ based on ensemble activity. Here, we used canonical correlation analysis (CCA) to assess fine-timescale coordination between *M1* and cerebellum. CCA has been recently employed in neuroscience studies to extract correlated population activity between two areas^39–43^. Specifically, CCA finds a linear combination of units in *M1* and cerebellum that represent maximally correlated activity across these areas (**Fig. 5a**). To establish that CCA of *M1* and cerebellar activity subspaces were significant, we compared the canonical variables of actual data with a distribution of canonical variable of trial-shuffled data (**Fig. 5b**; see methods). We used concatenated single-trial spiking activity binned at 50ms^40, 43^. The top component produced by CCA (known as CV1, canonical variable 1) is the axis of the *M1* and cerebellar subspaces that has the maximum correlation between the two areas (**Fig. 5c**). We first performed CCA between all *M1* task-related (*TR*) units (*M1 TR_d_* and *TR_i_* pooled together) and cerebellum *TR_i_* units. In order to have more M1 dimensions, we combined *M1 TR_d_* and *TR_i_* units as *M1 TR*. We found that this maximum correlation increased with neuroprosthetic learning for *M1 TR*–cerebellum *TR_i_* units. **Figure 5d** shows CCA changes in an example session from a single animal from *early* to *late* trials; and **Figure 5e** shows CCA change across all sessions for *M1 TR*– cerebellum *TR_i_* units (*early* canonical correlation: 0.30 ± 0.041, *late* canonical correlation 0.47 ± 0.038, mixed-effects model: *t(38)* = 3.96, *P* = 3.1 × 10^−4^). However, canonical correlation between *M1 TU*– cerebellum *TR_i_* units did not significantly increase across sessions (**Fig. 5f**; canonical correlation change: 0.21 ± 0.048 to 0.22 ± 0.051, mixed-effects model: *t(38)* = 0.38, *P* = 0.7; **Fig. 5g**).

**Figure 5.**
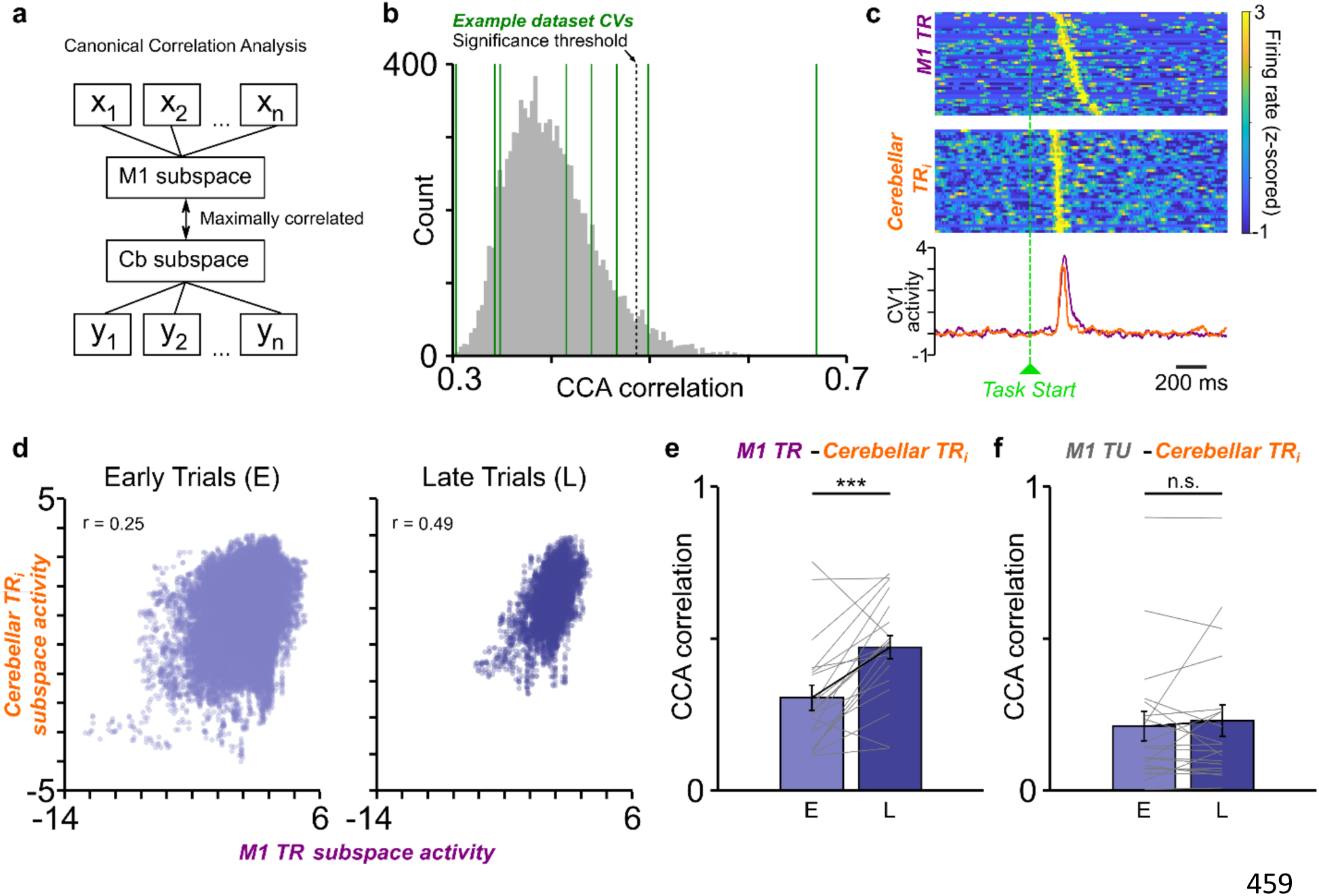
Increase in neural subspace correlation between task-related units of *M1* and cerebellum. ***a,*** Description of canonical correlation analysis (CCA). CCA finds a linear combination of binned spike counts from *M1* units (*x_1_*, *x_2_*, … *x_n_*) and cerebellum units (*y_1_*, *y_2_*, … *y_n_*) that maximizes the correlation between *M1* and cerebellum. ***b,*** Example identification of significant canonical variables (CVs, green lines) relative to trial-shuffled data (gray distribution, 10^4^ shuffles). Significant threshold at 95^th^ percentile of the distribution is shown in dotted gray line. Two canonical variables crossed this threshold in this example session. ***c,*** Single-trial *M1* task-related (*TR*) and cerebellar *TR_i_* spiking activity along with CV1 activation from *M1 TR*–cerebellar *TR_i_* CCA aligned to the task-start. ***d,*** *M1 TR* and cerebellum *TR_i_* subspace activity (from the CV1) around task-start (−2 to 2s) for an example session. Each dot represents one time bin of *early* or *late* trials from the session. Canonical correlation score is given by *r*. ***e,*** Change in the canonical correlation score from *early* to *late* trials across all sessions for *M1 TR* and cerebellar *TR_i_* units. ***f,*** Same as ***e*** for *M1 TU* and cerebellar *TR_i_* units. ***: p<0.001; n.s.: non-significant, p>0.05.

We also performed the CCA analysis for equal number of successful *early* and *late* trials (**Extended Fig. 5**). We found that the trend of increase in canonical correlation between *M1 TR*–cerebellum *TR_i_* units (**Extended Fig. 5a**; 0.33 ± 0.044 to 0.48 ± 0.04, mixed-effects model: *t(38)* = 3.13, *P* = 3.3 × 10^−3^) remained the same with this subset of trials. We also saw no significant change in the canonical correlation between *M1 TU*–cerebellum *TR_i_* units in these trials (**Extended Fig. 5b**; 0.24 ± 0.048 to 0.23 ± 0.043, mixed-effects model: *t(38)* = -0.05, *P* = 0.95).

Next, we performed this analysis for *M1 TR_d_*–cerebellum *TR_i_* units and *M1 TR_i_* –cerebellum *TR_i_* units. We found that the canonical correlation increased with neuroprosthetic learning even for *M1 TR_d_* –cerebellum *TR_i_* units. **Extended Fig. 5d** shows CCA changes in an example session for this pair from a single animal from *early* to *late* trials; **Extended Fig. 5e** shows CCA change across all sessions for *M1 TR_d_* –cerebellum *TR_i_* units (*early* canonical correlation: 0.14 ± 0.022, *late* canonical correlation 0.29 ± 0.033, mixed-effects model: *t(38)* = 4.30, *P* = 1.1 × 10^−4^). Moreover, we found that the subspace activity in the two structures became more precisely temporally correlated with learning as the higher the canonical correlation grew, the shorter the time to task-completion became. (**Extended Fig. 5f**). We observed that canonical correlation significantly increased from *early* to *late* trials even for *M1 TR_i_* –cerebellum *TR_i_* (**Extended Fig. 5g**; canonical correlation change: 0.31 ± 0.053 to 0.52 ± 0.042, mixed-effects model: *t(38)* = 4.51, *P* = 5.8 x 10^−5^).

### Cerebellum neural activity predicts M1 BMI-potent neural activity

With recent reports of *M1* activity being input driven during forelimb reaching behavior and secondary motor cortex’s (*M2*) modulatory influence over *M1* during *M1*–driven neuroprosthetic task^26, 40^, we wanted to check how the cerebellar *TR_i_* activity ‘integrated’ with *M1 TR_d_*, *TR_i_*, and *TU* activity. Upon a simple comparison of latency of *M1* and cerebellar spiking activities’ peaks, we observed that cerebellum *TR_i_* activity tended to peak before *M1 TR_d_* during *early* trials (time to peak for *M1* activity: 238.01 ± 6.42ms; and time to peak for cerebellum activity: 236.82 ± 6.48ms, mixed-effects model: *t(646)* = −0.07, *P* = 0.93) and *late* trials (time to peak for *M1* activity: 239.82 ± 8.26ms and time to peak for cerebellum activity: 211.16 ± 15.12ms, mixed-effects model: *t(646)* = −1.87, *P* = 0.06). This then lead us to develop a generalized linear model (GLM) to determine the relationship between cerebellum indirect activity and *M1* activity^44^.

We used cerebellum task-related indirect activity (*i.e.,* cerebellum *TR_i_* ’s) as a predictor of *M1* BMI–potent neural activity, where BMI–potent activity was *M1 TR_d_* activity or a ‘surrogate BMI–potent activity’ *for M1 TR_i_* ’s or *M1 TU* ’s which were used as the response variable for three different GLMs, namely, GLM–C*d* (Cerebellum *TR_i_* ’s è *M1 TR_d_* ’s), GLM–C*i* (Cerebellum *TR_i_* ’s è *M1 TR_i_* ’s) and GLM–C*U* (Cerebellum *TR_i_* ’s è *M1 TU* ’s), respectively (**Fig. 6a,b**). We also evaluated GLMs regression weights to analyze the temporal structure of these three predictions. For GLM-C*d*, we observed that various cerebellum *TR_i_* ’s exhibited their highest magnitude weight at different time lags across the population (**Fig. 6c**), indicating a broader timescale modulation of M1 *TR_d_* activity by cerebellum *TR_i_* ’s. Furthermore, numerous individual cerebellum units displayed large regression weights at multiple time lags, often encompassing both positive and negative weights (**Fig. 6d**). Similar regression weights timescales were observed for GLM– C*i* and GLM–C*U* models (**Extended Fig. 6**). However, we found that the *R*^2^ values for GLM–C*d* and GLM–C*i* models were significantly higher as compared to GLM–C*U* model (**Fig. 6e**; GLM–C*d*: *R*^2^ = 0.0504 ± 0.0225 to GLM–C*U*: *R*^2^ = –0.0358 ± 0.0302; mixed-effects model: *t(38)* = −2.40, *P* = 0.02; GLM–C*i*: *R*^2^ = 0.0617 ± 0.0233 to GLM–C*U*: *R*^2^ = –0.0358 ± 0.0302; mixed-effect model: *t(38)* = −2.69, *P* = 0.01). We did not find a significant difference between *R*^2^ values of GLM-C*d* and GLM-C*i* (mixed-effect models: *t(38)* = 0.42, *P* = 0.67).

**Figure 6.**
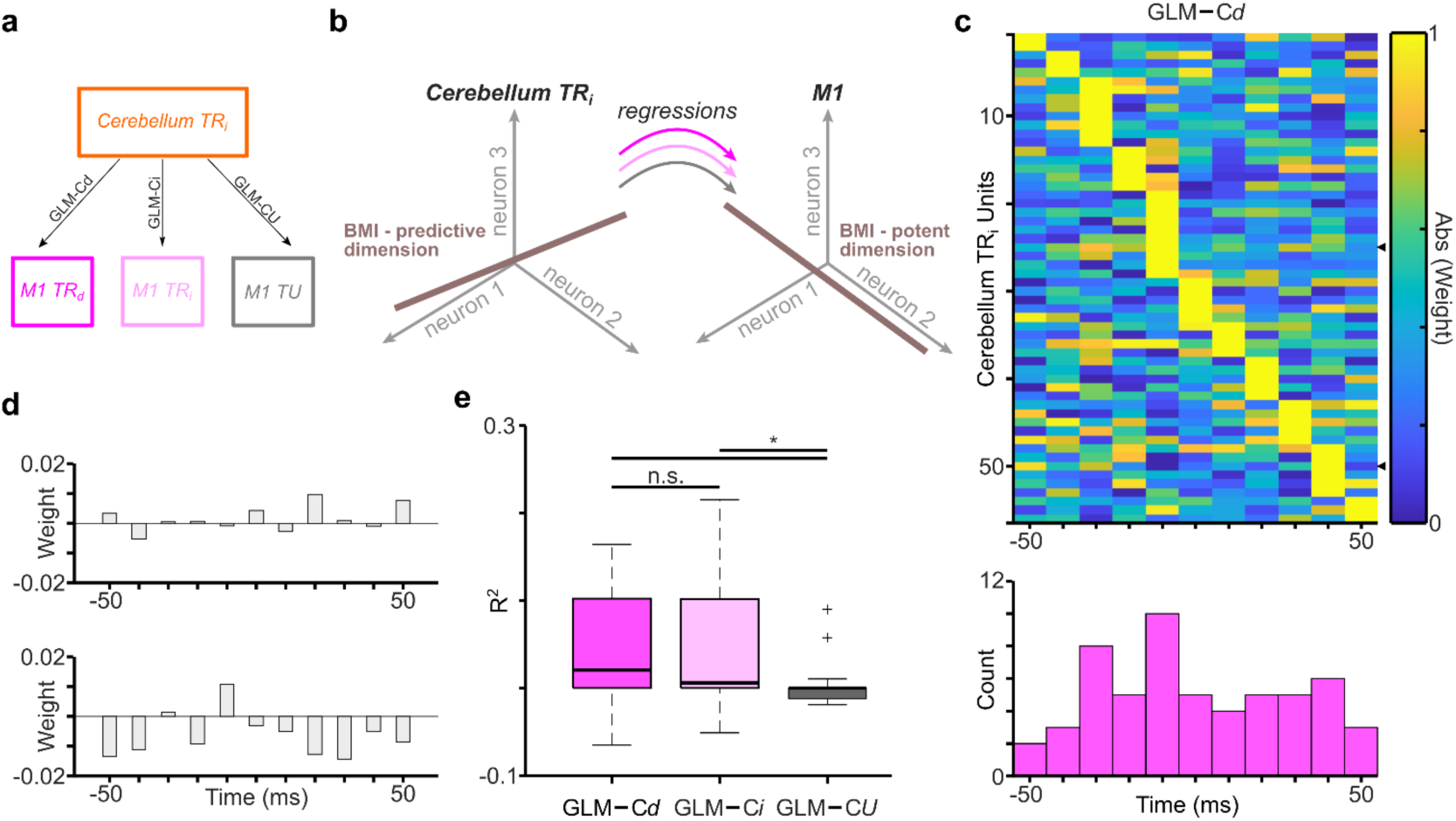
Cerebellum *TR_i_* neural activity predicts *M1* BMI-potent neural activity. ***a,*** Description of GLM model; GLM–C*d*: GLM model predicting *M1 TR_d_* activity from cerebellum *TR_i_* ’s; GLM–C*i*: GLM model predicting *M1 TR_i_* activity from cerebellum *TR_i_* ’s; GLM–C*U*: GLM model predicting *M1 TU* activity from cerebellum *TR_i_* ’s. ***b,*** Regression was used to identify a cerebellum neural population space that predicted BMI-potent *M1* activity. GLMs were fit to predict the *M1* BMI task-related direct (*TR_d_*)/ task related indirect *(TR_i_)*/ task unrelated (*TU*) neural state from cerebellum *TR_i_* activity; multiple time lagged copies of each cerebellum *TR_i_* unit were used as predictors. ***c,*** Distribution of regression weight magnitude in one example session for GLM–C*d* model (fitted to neural data binned at 10ms). *Top*, for each cerebellum *TR_i_* unit, regression weights were assigned for a variety of time lags. To emphasize the time of the maximum absolute weight of each neuron, values here are normalized to each neuron’s maximum value, and the absolute value of those weights is indicated by color. Units are sorted according to the time of the largest magnitude weight. Tick marks on the right edge indicate the units shown in ***d***. *Bottom*, histogram of the *τ* values with the largest magnitude weight for this dataset. ***d,*** Example non normalized weights for two cerebellum *TR_i_* neurons from one example session (neural data binned at 10 ms). Height of bars indicates weights for example neurons at different time lags (*τ*) relative to the *M1* BMI–potent activity, with negative *τ* values meaning that cerebellum *TR_i_* leads. ***e,*** Box–plot comparing *R*^2^ values for 3 different GLM models (fitted to neural data binned at 50 ms), GLM–C*d*, GLM–C*i* & GLM– C*U*; left to right (box–plot conventions are same as Fig. 3c). *p < 0.05; n.s. non-significant; p > 0.05.

### Optogenetic inhibition of cerebellum cortex and nuclei impairs neuroprosthetic performance

Next, we wanted to assess the necessity of cerebellar activation for *M1*–driven neuroprosthetic learning and hence, we performed optogenetic inhibition of cerebellar cortex and deep cerebellar nuclei (*DCN*) on neuroprosthetic skill learning. We used a red-light shifted halorhodopsin–JAWS for inhibiting neural activity (see Methods). First, we found that JAWS was robustly expressed in cerebellar cortical and *DCN* neurons (**Fig. 7 and 8**). When we looked at the activity of cerebellar cortical neurons under optical illumination acutely, we found that JAWS activation led to strong inhibition of these neurons (**Extended Fig. 7a,b**). Optogenetic inhibition significantly reduced firing across cerebellar cortical neurons (**Extended Fig. 7c**; *Stim_pre_*: 27.30 ± 2.88Hz, *Stim_ON_*: 4.99 ± 0.67Hz and *Stim_post_*: 20.16 ± 2.09Hz; *Stim_pre_* vs *Stim_ON_* mixed-effects model: *t(88)* = −7.69, *P* = 1.9 × 10^−1^^1^; *Stim_post_* vs *Stim_ON_* mixed-effects model: *t(88)* = 7.05, *P* = 3.8 × 10^−10^), with a reduction in 84.44% of recorded cells during *Stim_ON_* (*n = 45*).

**Figure 7.**
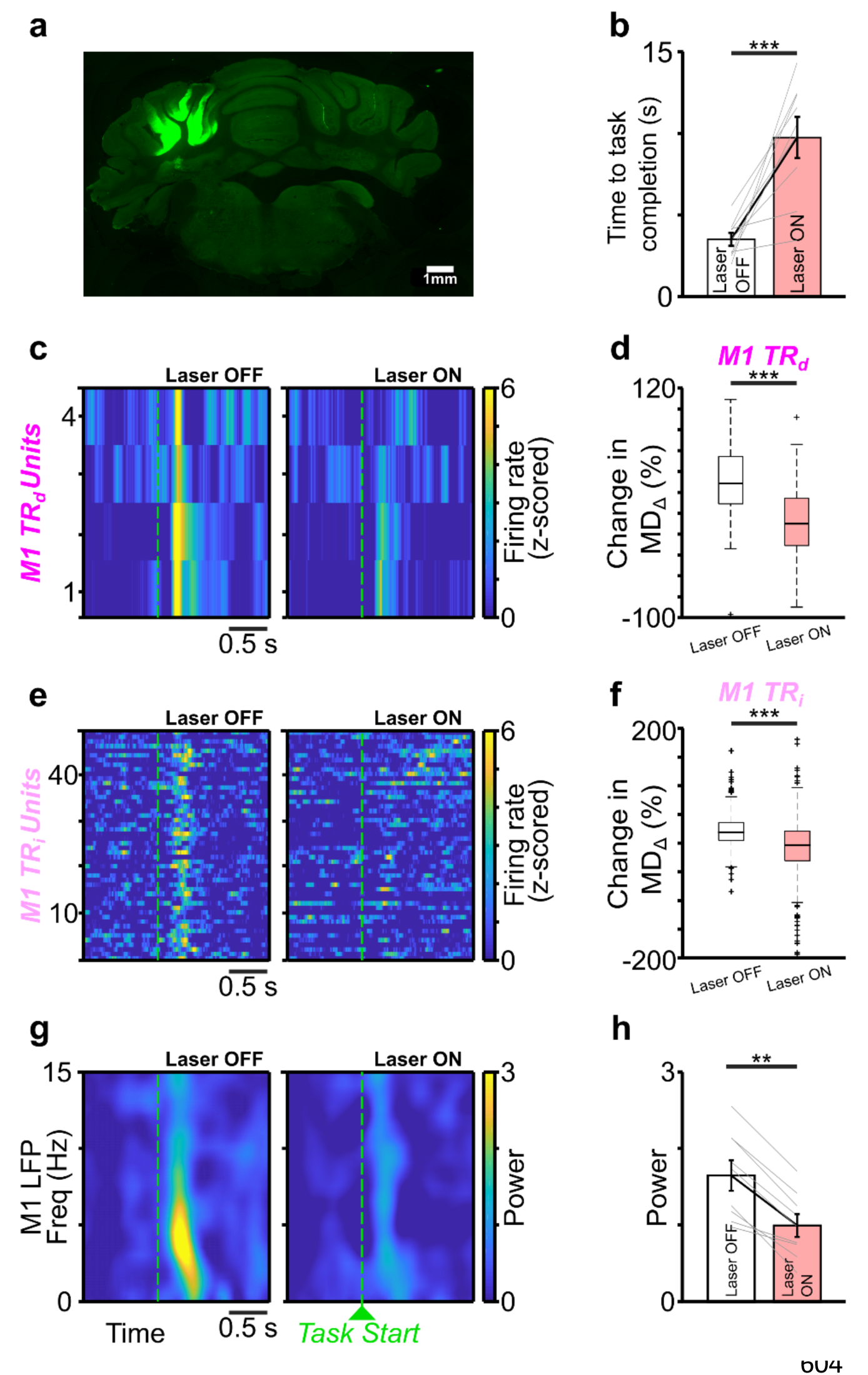
BMI performance gets impaired with cerebellar cortical inhibition. ***a,*** Fluorescence image of a coronal brain section showing neurons expressing JAWS (green) in the cerebellar cortex (Simplex and Crus I). ***b,*** Cerebellar cortical inhibition increases time to task completion. ***c,*** PETH of example *M1 TR_d_* units from different sessions, during *late* trials, with (Laser ON) and without (Laser OFF) cerebellar inhibition. ***d,*** Box-plot showing change in modulation depth (*MD_Δ_*) of *M1 TR_d_* units from *early* to *late* trials, with and without cerebellar inhibition (box–plot conventions are same as Fig. 3c). ***e,*** Same as ***c*** for *M1 TR_i_* units. ***f,*** Same as ***d*** for *M1 TR_i_* units. ***g,*** Spectrograms of an example *M1* LFP channel showing an absence of 3–6 Hz power during cerebellar inhibition (*left*) in *late* trials. The 3–6 Hz power emerges during *late* trials in the same day session where cerebellar cortical inhibition was suspended. ***h,*** 3–6 Hz power emerges during *late* trials on the same day session where cerebellar inhibition was not done. ***p<0.001. **p<0.01.

**Figure 8.**
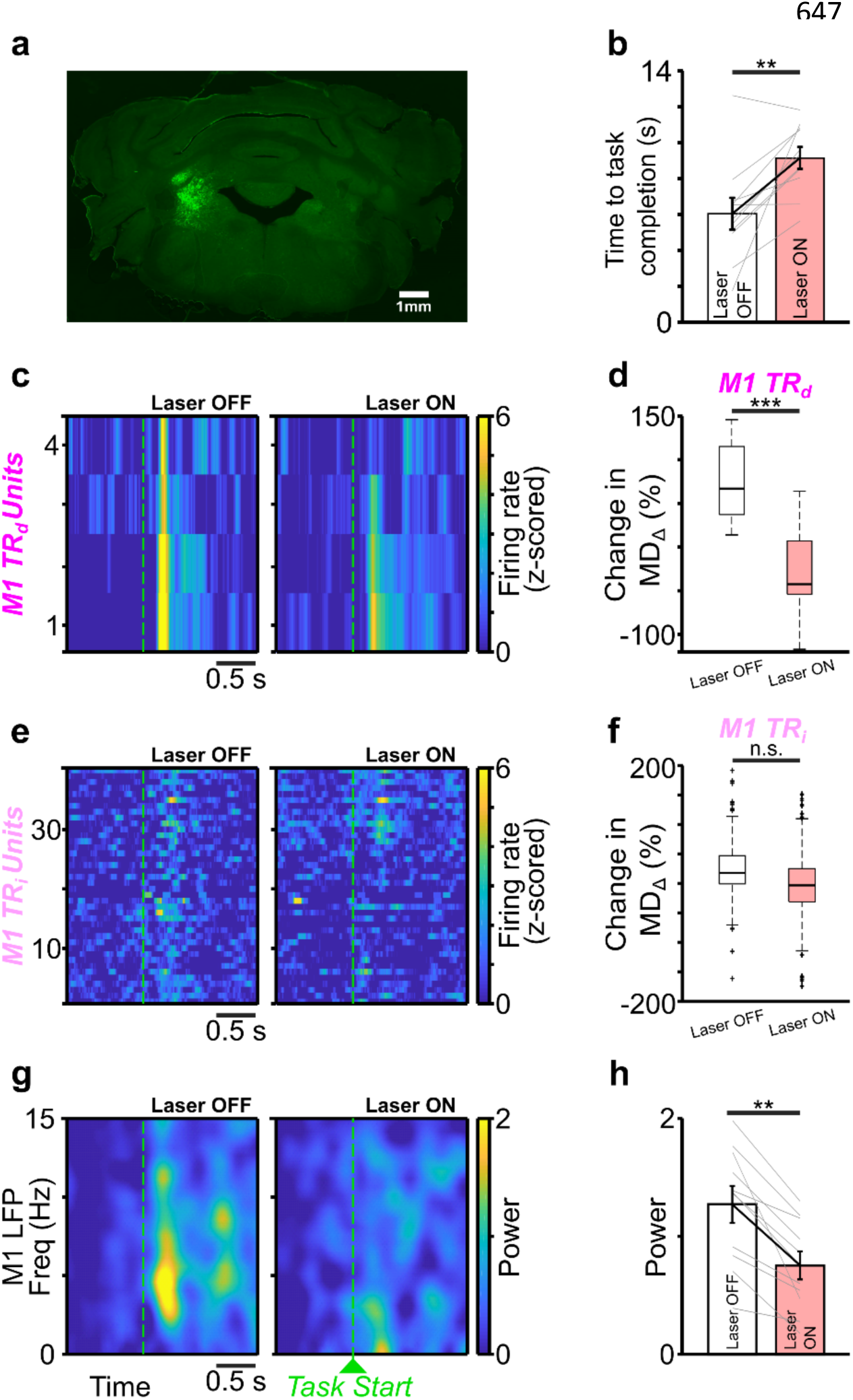
BMI performance gets impaired with *DCN* inhibition. ***a,*** Fluorescence image of a coronal brain section showing neurons expressing JAWS (green) in the deep cerebellar nuclei (*DCN*). ***b,*** *DCN* inhibition increases time to task completion. ***c,*** PETH of example *M1 TR_d_* units from different sessions, during *late* trials, with (Laser ON) and without (Laser OFF) *DCN* inhibition. ***d,*** Box–plot showing change in modulation depth (*MD_Δ_*) of *M1 TR_d_* units from *early* to *late* trials, with and without *DCN* inhibition (box plot conventions are same as **Fig. 3c**). ***e,*** Same as ***c*** for *M1 TR_i_* units. ***f,*** Same as ***d*** for *M1 TR_i_* units. ***g,*** Spectrograms of an example *M1* LFP channel showing an absence of 3–6 Hz power during *DCN* inhibition (*left*) in *late* trials. The 3–6 Hz power emerges during *late* trials in the same day session where *DCN* inhibition was not done. ***h,*** Bar-plot showing that 3–6 Hz power emerges during *late* trials on the same day session where *DCN* inhibition was not done. ***p<0.001. *p<0.05

We first performed optogenetic inhibition of the cerebellar cortex in chronically implanted rats during the BMI task training (see Methods). When we inhibited the cerebellar cortex in rats that had already gained proficiency in neuroprosthetic task performance (i.e., during *late* trials), we found that time to successful completion of neuroprosthetic task increased (**Fig. 7b**; 3.50 ± 0.39s without cerebellum cortex inhibition and 9.75 ± 1.26 s with cerebellum cortex inhibition, mixed-effects model: *t(16)* = 5.28, *P* = 7.3 × 10^−5^). We also found that the distribution of time to task completion for all trials across rats was significantly different without cerebellum cortex inhibition as compared to with cerebellum cortex inhibition **(Extended Fig. 8a**; Kolmogorov–Smirnov two-sample test: *P* = 3.4 × 10^−44^).

Furthermore, we also found that cerebellar cortex inhibition showed a decrease in the firing rate of *M1 TR_d_* units (**Fig. 7c** & **Extended Fig. 8b**). This led to a decrease in the *MD_Δ_* of *M1 TR_d_* units during trials when cerebellum cortex was optogenetically inhibited (**Fig. 7d**; 46.70 ± 8.6% (mean ± s.e.m.) without cerebellum cortex inhibition and –23.89 ± 9.90% with cerebellum cortex inhibition, mixed-effects model: *t(93)* = −3.85, *P* = 2.1 × 10−4). We also found that cerebellar cortex inhibition affected *M1 TR_i_* units at a population level (**Fig. 7e,f**; *MD_Δ_* for *M1 TR_i_* units: 29.53 ± 4.29% (mean ± s.e.m.) without cerebellum cortex inhibition and –5.11 ± 2.67% with cerebellum cortex inhibition, mixed-effects model: *t(821)* = – 4.99, *P* = 7.16 × 10^−7^). Cerebellar cortical inhibition also impacted *M1* 3−6 Hz LFP power. During *late* trials, 3−6 Hz *M1* power was reduced under cerebellar inhibition (**Fig. 7g,h**; z–scored *M1* power without cerebellum cortex inhibition: 1.64 ± 0.19; and *M1* power with cerebellum cortex inhibition: 0.99 ± 0.14; mixed effects model: *t(16)* = −2.95, *P* = 9.3 × 10^−3^).

While these experiments spoke about loss of performance on the neuroprosthetic task when cerebellar cortex was inhibited, we also analyzed neuroprosthetic task performance vis-à-vis the order of cerebellar cortex inhibition (*i.e.*, the effects of cerebellar cortical inhibition in either first BMI block, *BMI_1_*, versus the second BMI block, *BMI_2_*, within a day with the same *M1 TR_d_* units; see Methods for details on the order of two BMI blocks within a day). We found that neural processing in the cerebellar cortex impaired *M1*– driven neuroprosthetic control irrespective of the order of inhibition, this impairment was significantly more pronounced when inhibition occurred in *BMI_1_* (**Extended Fig. 8d**; time to task completion on *late* trials without cerebellum cortex inhibition in *BMI_1_*: 4.26 ± 0.77s versus time to task completion on *late* trials with cerebellum cortex inhibition in *BMI_2_*: 6.77 ± 2.17s, mixed-effects model: *t(6)* = 1.44, *P* = 0.19. **Extended Fig. 8e**; time to task completion with cerebellum cortical inhibition in *BMI_1_*: 11.91 ± 0.87s versus time to task completion without cerebellum cortical inhibition in *BMI_2_*: 3.10 ± 0.50s, mixed-effects model: *t(8)* = –10.89, *P* = 4.4 × 10^−6^).

When we performed the optogenetic inhibition of the *DCN*, we observed similar deficits in neuroprosthetic control and *M1* physiology. We found that time to successful completion of neuroprosthetic task increased (**Fig. 8b**; without *DCN* inhibition: 6.04 ± 0.88s; and with *DCN* inhibition: 9.15 ± 0.60s; mixed-effects model: *t(20)* = 3.19, *P* = 4.6 × 10^−3^). We also found that the distribution of time to task completion for all trials across rats was significantly different between the two conditions (**Extended Fig. 9a**; Kolmogorov– Smirnov two-sample test: *P* = 2.2 × 10^−17^). We found that *DCN* inhibition showed a decrease in the firing rate of *M1 TR_d_* units (**Fig. 8c** & **Extended Fig. 9b**). This led to a decrease in the *MD_Δ_* of *M1 TR_d_* units during trials when *DCN* was optogenetically inhibited (**Fig. 8d**; without *DCN* inhibition: 108.89 ± 17.95% (mean ± s.e.m.); and with *DCN* inhibition: −41.76 ± 10.17%; mixed-effects model: *t(51)* = −3.85, *P* = 1 × 10−6). *DCN* inhibition also showed non-significant reduction in *MD_Δ_* of *M1 TR_i_* units at a population level (**Fig. 8e,f**; *MD_Δ_* for *M1 TR_i_* units without *DCN* inhibition: 39.15 ± 5.13%; and with *DCN* inhibition : 11.77 ± 15.04%, mixed-effects model: *t(610)* = –1.07, *P* = 0.28). We found that 3–6 Hz M1 LFP power was significantly reduced with *DCN* inhibition as well during *late* trials (**Fig. 8g,h**; *z–scored M1* power without *DCN* inhibition: 1.26 ± 0.15; and *M1* power with *DCN* inhibition: 0.75 ± 0.11; mixed effects model: *t(20)* = −3.20, *P* = 4.4 × 10^−3^).

As with cerebellar cortical inhibition, we performed *DCN* inhibition either in *BMI_1_* or *BMI_2_* and assessed neuroprosthetic task performance. Here, we observed that more pronounced neuroprosthetic task impairments occurred when *DCN* was inhibited in *BMI_2_* (**Extended Fig. 9d**; time to task completion in *late* trials without *DCN* inhibition in *BMI_1_*: 4.88 ± 1.26s versus time to task completion with *DCN* inhibition in *BMI_2_*: 9.36 ± 1.14s, mixed-effects model: *t(8)* = 3.27, *P* = 0.01; **Extended Fig. 9e**; time to task completion in *late* trials with *DCN* inhibition in *BMI_1_*: 8.98 ± 0.78 s versus time to task completion in *late* trials without *DCN* inhibition in *BMI_2_*: 7.01 ± 1.26 s, mixed-effects model: *t(10)* = −1.58, *P* = 0.14). Hence, while we observed significant neuroprosthetic task impairments with cerebellar inhibition both at the level of the cortex and its deep nuclei when we analyzed both *BMI_1_* and *BMI_2_* together, but upon parsing these two BMI blocks, we observed more significant impairment of cerebellar cortical inhibition on *BMI_1_* and *DCN* inhibition on *BMI_2_*.

## Discussion

In this study, we found an emergent 3–6 Hz activity in the *M1* and cerebellum LFPs and task-related direct and task-related indirect spiking in these regions was coordinated with this activity, but not the task unrelated *M1* spiking activity. Additionally, we found that neuroprosthetic task learning led to increased correlated neural subspace activity between *M1 TR_d_* –cerebellar *TR_i_* units and *M1 TR_i_* –cerebellar *TR_i_* units, but not for *M1 TU* –cerebellar *TR_i_* units. Furthermore, we found that cerebellar *TR_i_* activity well predicted *M1 TR_d_* and *M1 TR_i_* activity but not *M1* TU activity. Finally, we found that optogenetic inhibition of the cerebellum, either in the cerebellar cortex or its deep nuclei led to neuroprosthetic task performance impairments (as indicated by increased time to task completion) and weakening of *M1* task-related activity. These findings suggest that cerebellum plays an important role by providing influence on *M1* neural activity that is related to neuroprosthetic task output. Furthermore, we found that cerebellar task related indirect activity developed a preferential relationship with task-related *M1* direct and indirect activity suggesting that cerebellum *TR_i_* ’s had a more privileged relationship with task-relevant neurons of *M1* (*TR_d_* ’s and *TR_i_* ’s). Although our GLMs suggested that cerebellum *TR_i_* ’s were predictive of both *M1 TR_d_* ’s and *TR_i_* ’s activity, our optogenetic experiments showed that *M1 TR_d_* ’s and *TR_i_* ’s were impacted by cerebellar inactivation. This might be indicative of cerebellum processing to be linked to a preferential coordination of *M1* task-relevant units. Our elaboration of the cerebellum’s role in *M1*–driven neuroprosthetic motor control is consistent with the cerebellum’s role in fine-tuning movement, as well as co-emergent activity that has been reported in these areas with learning new motor skills^2, 25^. This study helps elucidate cerebellar contributions to *M1*–driven neuroprosthetic control and can help improve BMI functionality in the future. For example, BMI paradigms could incorporate cerebellar indirect signals for improving BMI controllers.

### Emergent mesoscopic dynamics across M1-cerebellum

One of our first findings was an emergence of 3–6 Hz coherence in *M1*–cerebellum LFPs associated with learning, and neurons in both these regions also showed enhanced phase-locking to this oscillation during task-relevant periods as revealed through STA. Similar observations have been made in a neuroprosthetic study that looked at task-related cells in cortico-striatal networks^20^. Such coherence can serve to enhance communication during task period between task-relevant cell populations across the larger motor networks that should integrate signals for optimal behavioral output. Such coherent activity may allow for flexible use of task-relevant cells in either region. The synchrony that we observed in 3– 6Hz band is consistent with other work that has showed low-frequency coherence between *M1* and other motor regions during learning^6, 7^. One of the possibilities for the 3–6 Hz increased coherence that we observed could be a result of common neural-drive to both these regions (or one of the regions driving the other). It is also pertinent to mention that similar low-frequency oscillations in the neocortex can be used to decode reach-related activity and predict spiking phase across multiple behavioral states^45, 46^. Such activity is also correlated with multiphasic muscle activations and timing of movements during motor tasks^46–49^. Recent work also suggests that oscillatory dynamics reflect an underlying dynamical system^48^. This previous work argues that this low-frequency activity represents an intrinsic property of motor circuits associated with precise motor control. Our findings extend this body of work by showing similar low frequency dynamics in both *M1* and cerebellum cortex (**Fig. 2**). The exact origin of these oscillations and underlying generators remains unknown. While such oscillations were shown to involve striatum in rodent reaching task^6^ or thalamocortical activity^50^, so far, our results here raise the possibility of cerebellar involvement. Further work can probe interactions between *M1* and the broader motor network to pinpoint the drivers of the electrophysiologic changes seen during learning this skill.

### Using BMIs to study cross-region coordination in motor control

BMI offers investigations of connected regions by selecting target neural activity (that dictate output) in one region and concurrent examination of another region as task performance improves. Implementing this strategy, we aimed to disentangle cerebello-cortical communication as *M1* direct control was learned. Importantly, both *M1* and cerebellum have direct connections to the spinal cord^8–10^, and are implicated in movement control^3, 4, 6^. In our BMI paradigm, we randomly selected target neural activity in *M1* (enforced by the decoder), which is unlikely to be correlated with processes in other brain regions. This permitted us to examine how cross-area communication during BMI control facilitates control. *M1* and cerebellum are reciprocally connected^1, 2^, and while some extent of coupling of task-related activity in *M1* and cerebellum is not surprising, it is unknown how the task-related indirect cerebellum activity interacts with behavioral output-specific direct activity versus task-related indirect activity or task-unrelated activity of *M1* neurons. It is important to note that we didn’t find efficient neuroprosthetic control to be linked to limb movements or other idiosyncratic movements (**Extended Fig. 2**).

One of the striking findings of our work was that canonical correlation increased between *M1* and cerebellum task-related units, but not for *M1 TU*–cerebellum *TR_i_* units (**Fig.5 & Extended Fig. 5**). Similarly, 3–6 Hz oscillatory activity modulated task-related direct and indirect activity in *M1* as well as cerebellar indirect activity, but not *M1 TU* activity (**Figs. 3 and 4**). Our predictive model corroborated this finding and found that cerebellar *TR_i_* activity predicted *M1 TR_d_* and *TR_i_*, but not *M1 TU* activity. These analyses revealed that the task-related cerebellar cortical activation communicated more strongly with task-relevant pools of *M1* activity. These ideas are consistent with theoretical studies that suggest that cerebellar cortex helps in task-relevant dimensionality expansion that might aid in learning^51^.

### Roles of multiplexed cross-area interactions

Motor control involves signals at longer time scales appropriately integrating with shorter time scales across several spatially segregated regions to deliver movement precision^52^. Little is known about how these signals at varying spatiotemporal scales are interacting during motor control. Simultaneous recordings are best suited to understand these interactions^6, 39, 53–55^. Further, a majority of studies that have used simultaneous recordings in motor regions use extensively trained animals performing natural motor tasks^54–57^. Understanding *M1*–cerebellum interactions in such context raises important concerns: *(i)* since both *M1* and cerebellum are directly controlling movement, it is difficult to ascertain modulatory influence of one area over the other; and *(ii)* overtrained animals may have transitioned to an *‘automatic’* state^58^, and it may no longer be suitable to investigate emergent dynamics in interacting structures. These confounds limit the inferences made about *M1*–cerebellum interactions in experiments with extensively trained animals.

Here we have shown that in an *M1*–driven *BMI* task, cerebellum neural processing was crucial. While *BMI* performance was impacted due to cerebellar inhibition overall, we found that this performance was impacted to varying degrees by cortical versus *DCN* inhibition based on the order of inhibition. Cerebellar cortical inhibition impacted *BMI_1_* task performance significantly (**Extended Fig. 8e**), which is indicative of cerebellar cortical processing having an instructional role early-on through the olivo-cerebellar system, consistent with the notion of cerebellar role in skill acquisition^59^. Our optogenetic inhibition may have disturbed cerebellar cortical processing by either altering inhibitory inputs onto the Purkinje cells (*PCs*) from stellate cells, basket cells, or other molecular layer interneurons (*MLIs*), or the granule cells’ parallel fiber (*PF*) inputs to *PCs* or *PCs* themselves. When we inhibited *DCN*, we found that *BMI_2_* task performance was significantly impaired (**Extended Fig. 9d**) when the control was already well learned. This is consistent with the role of cerebellar output in fine-tuning ongoing movements^24, 25^. Future work with cell-specific inhibition in the cerebellar cortex or *DCN* can test the effects on the neuroprosthetic task performance. Overall, we showed the involvement of the cerebellum in *M1*–driven neuroprosthetic control, which has not been shown before.

Cerebellar involvement in *M1*–driven *BMI* task is further cemented by the fact that cerebellar *TR_i_* activity had strong modulation in *late BMI* trials as well (**Fig. 1i**), indicative of an ongoing modulatory influence over *M1* to sustain proficiency in the task. Our canonical correlations (**Fig. 5**), spike–LFP coordination (**Fig. 3,4**), and predictive model (**Fig. 6**), all showed that task-related pools of *M1* and cerebellum develop a preferential relationship. This, with other recent work, leads us to conclude that *M1* task-relevant cells multiplex signals locally^11, 44^ as well as from distant-area cerebellar activity^60^. Our regression of *M1* BMI potent space activity also indicated that cerebellum *TR_i_* units showed a broader timescale influence of cerebellum on *M1 TR_d_* activity (this is different from the shorter time scale influences seen between *M1 TR_d_* and *M1 TR_i_* units^44^). Such broader influence of cerebellar activity may also be related to coordination in the larger motor network. Larger network activity may represent attention regulation^61^, motivation^62^, or coordination of the motor task with sensory feedback^63^.

### Neural dynamics over the course of BMI learning

Natural motor learning is known to involve an early phase marked exploration and high variability with a transition to late stage when the skill is consolidated^64–67^. Our paradigm here is focused on early exploratory *BMI* learning by using mostly single sessions (within a day). *BMI* studies that allow for sleep consolidation^11, 12, 18^ or use multiple days of learning^15, 68^ find that *M1 TR_i_* units weaken their modulation through the course of extensive training. Future work can test if cerebellar activation aided in such credit assignment as some of the optogenetic effects were selective for *M1 TR_d_* ’s in our work. It is also possible that cerebellar *TR_i_* ’s may exhibit similar weakening (as *M1 TR_i_* ’s) over time. However, local versus cross area interactions may differ in the long-term. One recent study had focused on cross-area activity through multiple days of training, and they found that indirect task-related modulation persists in several cortical areas^23^. Work that has looked at extensively trained mice on motor task has shown sustained activity in the cerebellum^25^. There is further evidence from studies of natural learning that emergent activity in cortico-cerebellar networks becomes stronger as task proficiency increases^7^.

To summarize, our studies leveraged a multiarea *BMI* paradigm to probe *M1*–cerebellar cross-area interactions. We demonstrated that oscillatory dynamics emerged as seen through LFPs across these regions that also modulated task-related spiking in both areas. Finer time scale analyses of spiking revealed that cerebellar *TR_i_* activity selectively influenced task-related artificial target activity within *M1*. This paradigm allowed us to manufacture an output-specific *M1* activation to examine internal motor networks dynamics, removing the constraints of movement performance. Thus, multiarea *BMIs*, through such impositions, allows for a more natural inside-out investigation of cross-area interactions in the motor network.

## Methods

### Animal preparation

Adult male Long-Evans rats were used in this study (*n = 15*, 300–500 g, 3 – 5 months old, Charles River Laboratories). All animal procedures were performed according to the protocol approved by the Institutional Animal Care and Use Committee at Cedars-Sinai Medical Center, Los Angeles. This ensured that the animals that were used in this research were acquired, cared for, housed, used, and disposed of in compliance with the applicable federal, state, and local laws and regulations, institutional policies and with international conventions to which the United States is a party. Animals were housed on a 14 h light and 10 h dark cycle (photoperiod is from 6 am to 8 pm) in a climate-controlled vivarium. Out of 15 rats, 7 were used in BMI with simultaneous *M1* and cerebellum recordings, 3 rats were used for BMI with cerebellar cortex optogenetic inhibition and 3 rats for *DCN* optogenetic inhibition. The remaining (*n = 2*) were used in acute cerebellar recording under optogenetic inhibition. Neural probes were implanted during a recovery surgery performed under isofluorane (1–3%) anesthesia. The analgesic regimen included the administration of 0.1 mg per kg body weight buprenorphine, and 5 mg per kg body weight carprofen. Post-operatively, rats were also administered 2 mg per kg body weight dexamethasone and 33 mg per kg body weight sulfatrim for 5 days. Ground and reference screws were implanted posterior to lambda contralateral to the recorded cerebellum, contralateral to the neural recordings. For *M1* recordings, 32– channel arrays (33-μm polyamide-coated tungsten microwire arrays) were lowered to a depth of ∼1,200–1,500 μm in either the left or right *M1*. These were implanted centered at 0.5 mm anterior and 3 mm lateral to the bregma^6, 7, 11, 12, 18, 69^. For cerebellar recordings we used 32 – 64 channel tetrodes (Neuronexus, MI) or shuttle-mounted polytrodes (Cambridge Neurophysiology, UK). The probes were lowered into the cerebellar cortex through a craniotomy centered at 12.5 mm posterior and 2.5-3 mm lateral to bregma. Shuttle mounted probes were moved across days and recorded from depths of 1.5-4 mm. Our target regions were Simplex/ Crus I and Crus II areas of the cerebellum*_7,70–72_*. Activity in these areas has shown modulation during upper limb motor behaviors in response to corticofugal fiber and forelimb stimulation, and during forelimb reaching task. We did not perform subject-specific implantation based on motor mapping. However, a subset of rats also performed reaching tasks and had robust activation during reaching^7^.

### Viral injections

We used a red-shifted halorhodopsin, Jaws (AAV8-hSyn-Jaws-KGC-GFP-ER2, UNC Viral Core), for neural silencing in 6 rats for optogenetic experiments^12, 18, 73^. Viral injections were done at least 3 weeks before chronic implant surgeries. Rats were anesthetized, as stated before and body temperature was maintained at 37 °C with a heating pad. Burr hole craniotomies were performed over injection sites, and the virus was injected using a Hamilton Syringe with 34G needle. 500-nl injections (100 nl per min) were made at two sites in the cerebellar cortex (11.5 mm posterior, 2.5 mm lateral to bregma and 11.5 mm posterior and 3.5 mm lateral to bregma; depth of 1–3 mm). In *DCN* as well, we performed viral injections at two sites (11.5 mm posterior, 2.5 mm lateral to bregma and 11.5 mm posterior and 3.5 mm lateral to bregma; depth of 6.1–6.3 mm). After the injections, the skin was sutured, and the animals were allowed to recover with same regimen as stated above. Viral expression was confirmed with fluorescence imaging.

### Electrophysiology

Units and LFP activity were recorded using a 128–channel TDT-RZ2 system (Tucker-Davis Technologies). Spike data were sampled at 24,414Hz and LFP data at 1,017.3Hz. ZIF (zero insertion force) clip-based digital head stages from TDT were used that interface the ZIF connector and the Intan RHD2000 chip which uses 192x gain. TDT’s RS4 data streamer was used to save all raw data at 24,414Hz in all animals except two, where only spike times and waveform snippets were saved. Only clearly identifiable units with good waveforms and high signal-to-noise were used. The remaining neural data was recorded for offline analysis. Behavior related timestamps (i.e. trial onset, trial completion) were sent to the RZ2 analog input channel using an Arduino digital board and synchronized to neural data.

We have used the term *‘unit’* to refer to the sorted spike recordings from both the MEA and silicon probe recordings. For both, we initially used an online sorting tool (Synapse, TDT) for neuroprosthetic control. We used waveform shape and the presence of an absolute/relative refractory period in the inter-spike interval (ISI) to judge quality of isolation. Specifically, a voltage-based threshold was set based on visual inspection for each channel that allowed for best separation between putative spikes and noise; typically, this threshold was at least 4 standard deviations (SD) away from the mean. Events were time-stamped and waveforms for each event were peak aligned. K-means clustering was then performed across the entire data matrix of waveforms. Automated sorting was performed by: *(1)* first over-clustering waveforms using a K-means algorithm (i.e., split into many mini-clusters), *(2)* followed by a calculation of interface energy (a nonlinear similarity metric that allows for an automated decision of whether mini-clusters are actually part of the same cluster), and *(3)* followed by aggregation of similar clusters. We conducted offline spike sorting in Plexon (where spike times and waveform snippets were saved) or Spyking Circus^7, 74^ (where spike data were saved at 24,414Hz).

### Behavior

After recovery, animals were typically acclimated for 1–2 day to a custom plexiglass behavioral box (**Fig. 1a**) prior to the start of experimental sessions. After acclimatization, rats were water restricted for BMI training. We monitored body weights daily to ensure that the weight did not drop below 95% of the initial weight.

Behavioral sessions were typically conducted for 1–2 hours. Recorded neural data was entered in real time to custom routines in Matlab (R2018b; Mathworks, Natick, MA). These then served as control signals for the angular velocity of the feeding tube. The rats performed ∼120 trials on average in a session. In a subset of sessions (*n = 13*), we also video-monitored the rat during the BMI training using a 30-fps camera (TDT RV2 video processor, USA).

### Neural control of the feeding tube

During the BMI training sessions, we typically selected one to four *M1* channels. The units on these channels (2–8) were assigned as *‘direct’* (*TR_d_*) units, and their neural activity was used to control the angular velocity of the feeding tube. If one channel was chosen for neuroprosthetic control, its neurons were associated with positive unit weight (*TR_d+_*) and if two or more channels were chosen, some channels neurons were associated with a positive unit weight (*TR_d+_*) and others with a negative unit weight (*TR_d–_*). We never assigned the units on the same channel with a positive and negative weights. These units maintained their stability throughout the recording as evidenced by stability of waveform shape and interspike-interval histograms (**Extended Fig. 10**). We binned the spiking activity into 50ms bins. We then established a mean firing rate for each neuron over a 3–5 minutes of baseline period. The mean firing rate was then subtracted from its current firing rate at all time points.

The specific transform that we used was:

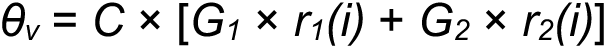

where *θ_v_* was the angular velocity of the feeding tube, *r_1_(i)* and *r_2_(i)* were firing rates of the direct units (*TR_d+_* and *TR_d–_*, respectively). *G_1_* and *G_2_* were fixed unit weights, *i.e.*, +1 and −1, respectively. *C* was a fixed constant (Gain) that scaled the firing rates to angular velocity. The animals were then allowed to control the feeding tube via modulation of neural activity. The tube started at the same position at the start of each trial (*P_1_* in **Fig. 1a**). The calculated angular velocity was added to the previous angular position at each time step (50 ms). During a trial, the angular position that was controlled from the *TR_d_* activity had the limits of 0° (*P_1_*) to 45° (*P_2_*). If the tube was controlled successfully to the *‘target position’* (*P_2_* in **Fig. 1a**), then a water reward was delivered at the final resting position set to 62°. In the beginning of a session, most rats were unsuccessful at bringing the feeding tube to final target position *P_2_*. Rats steadily improved control and reduced the time to completion of the task (*i.e.*, moving the tube from position *P_1_* to position *P_2_*) during a session. Multiple learning sessions were obtained from each animal. Consistent with past studies, we found that incorporation of new units into the control scheme required new learning^11, 75–77^. While we did not check whether the *TR_d_* units that we selected were part of the manifold^14^, but upon checking the covariance between *M1 TR_d_* decoder neurons during intertrial periods, we did not find a significant change with learning (*early* to *late* trials: 64.90 ± 29.60 to 108.06 ± 59.31; mixed-effect model: *t*(38) = 0.68, *P* = 0.49). This indicates that *TR_d_* units we selected were likely from the same manifold. We also found that the covariance between *M1 TR_d_* and *M1 TR_i_* units did not change with learning (*early* to *late* trials; 89.43 ± 33.14 to 163.38 ± 92.71; mixed-effect model: *t*(38) = 0.96 *P* = 0.34) indicating the *M1 TR_d_* and *M1 TR_i_* units also likely belonged to the same circuit.

### Optogenetics

Optogenetic experiments were carried out in JAWS injected rats using a high-power laser (50 mW/mm^2^: Laserglow Technologies, USA) emitting a 625 nm beam. A subset of rats (*n = 3*) was implanted with a 200 μm diameter optic fiber cannula (Doric Lenses) over Crus I/Crus II region of the cerebellar cortex and a 32-channel microwire array (TDT Florida) was implanted in *M1*. In another set of rats (*n = 3*) we implanted a cannula in *DCN* and a 32–channel microwire array in *M1*. During the behavior experiments, the laser was turned on every 50ms at 40% duty cycle to inhibit cerebellar activity after the trial onset for the total duration of the trial (15s). We performed two sessions each day with ∼100 trials in each session. Each day, we alternated between turning the laser on in the first session or the second session (**Extended Fig. 8c & Extended Fig. 9c**). Another subset of rats (*n = 2*) was implanted with a fiber-optic cannula mounted on a silicon probe (Cambridge Neurotech, UK) in the cerebellum, and the recordings were performed under isoflurane anesthesia. In every trial, we recorded 5s of baseline followed by 5s for which the laser was on and another 5s thereafter with laser off. We performed 30 such repetitions.

### Histology

After the experiment, rats were deeply anaesthetized with isoflurane (4–5%), then exsanguinated and perfused with 4% paraformaldehyde (PFA). The brains were extracted and stored in 4% PFA for up to 72 hours. The brains were then transferred to a solution of 30% sucrose and stored for sectioning. We performed sagittal or coronal sections of the brain using a cryostat (Leica, Germany) and stored them in phosphate buffered saline for imaging. Images were mounted on slices and imaged using a microscope (Keyence, Japan). The location and depth of the silicon probe in the brain were traced by *DiI* depositing on the electrodes prior to their implantation and by looking afterwards at the fluorescent dye present in the histological slices (**Extended Fig. 11**). The expression of JAWS virus was imaged in coronal sections of the cerebellum (**Fig. 7a** and **Fig. 8a**).

### Data Analysis

#### Sessions and changes in performance

Offline analyses were performed in MATLAB (R2020b) with custom-written routines. A total of 20 training sessions recorded from 7 rats were used for our initial analysis. In addition, we analyzed 18 separate sessions across 3 rats where optogenetic inhibition of the cerebellum cortex was performed, and 12 sessions across 3 rats where optogenetic inhibition of *DCN* was done. For **Fig. 1c,d** we compared changes in task performance across sessions. Specifically, we compared the performance change by calculating the mean and standard error of the mean (s.e.m) of the time to completion during the first and last 30% of trials (referred as *early* and *late* trials respectively). Furthermore, we also compared the performance in *early* and *late* trials by calculating the percentage of unsuccessful trials.

#### Task-related activity

The distinction between *TR_d_*, *TR_i_,* and *TU* units was based on the significant modulation over baseline firing activity of a unit after trial onset (i.e., peak of modulation at the time > 2.5 SD above the baseline period). We called this the modulation depth (*MD_Δ_*) of each unit. We took the difference between this modulation from *early* to *late* trials to compute the change in *MD_Δ_* for each *TR_d_* and *TR_i_* unit (**Fig. 7d,f** & **Fig. 8d,f**).

#### LFP analysis

Artifact rejection was first performed on LFP signals to remove broken channels and noisy trials. LFPs were then z-scored, median referenced and evoked-activity was subtracted separately for *M1* and cerebellum. LFP power was calculated on a trial-by-trial basis and then averaged across channels and animals, with wavelet decomposition using the EEGLAB function *newtimef*^78^. *M1*–cerebellum LFP coherence was calculated for each pair of channels using the EEGLAB function *newcrossf*^78^. All the comparisons were done between *early* and *late* trials. For this analysis, across all the early trials, only trials where time to task completion was over 10s were included. Across all the *late* trials, only trials where time to task completion was under 5s were included.

We also performed LFP power and coherence comparisons between the equal number of successful trials from *early* and *late* learning. From the *early* phase we included successful trials which were over 10s, and from the *late* phase we included the equal number of trials where time to task completion was under 5s. We performed LFP power and coherence analysis using the same EEGLAB functions that are mentioned above.

#### Spike-triggered averaging (STA)

We calculated the STA to measure how spikes locked to the 3–6 Hz LFP oscillations, both in the *M1* and the cerebellum. We used filtered (3–6Hz) median LFP from each region for this analysis. For filtering, we used the EEGLAB function *eegfilt*. We used the first 4s after the start of the trial to calculate these STAs. For every unit, we concatenated the spikes from *early* trials in a spike vector and from *late* trials in another vector. Before STA calculation, we equaled the length these vectors. Then, we extracted 2s of LFP around every spike-time in those vectors and average it to get *early* and *late* STAs for a given unit. To calculate the change in modulation for every unit, we looked at the difference between the minimum and maximum peak in a 300ms window around a spike in the averaged STA of *early* and *late* trails and then calculated this change from *early* to *late* trials in percentage. We also calculated STA during inter-trial interval. Here, we used from 2s to 6s after the end of the trial and applied the same steps to compute the STA as described above. Furthermore, we repeated the STA analysis for the task-period and the inter trial interval by using the equal number of successful only trials from *early* and *late* learning.

#### Spike-LFP phase analysis

To study the phase relationship between spiking and LFP activity, we generated histograms of the LFP phases at which each spike occurred for a single unit to LFP channels that showed an increase in power from *early* to *late* trials, in a 2.45s window around task-start (–0.25 s before to 2.2 s after movement onset) across all trials of a session (**Fig. 4a,b**). The LFP channels were filtered in the 3–6 Hz band. All units were compared with the same selected *M1* and cerebellum LFP channels from *early* to *late* trials. The histograms were generated for each unit–LFP channel pair both within and across regions. For every pair, we then calculated the Rayleigh’s z–statistic for circular nonuniformity. These z–statistics were then used to calculate the percentage of significantly nonuniform distributions across unit–LFP pairs with a significance threshold of *P* = 0.05 (**Fig. 4c,d,e,f**). A significantly nonuniform distribution signifies phase preference for spikes of a unit to an LFP signal.

#### Canonical correlation analysis

We identified shared cross-area subspaces between *M1* and cerebellum using canonical correlation analysis (CCA). This method identifies the maximally correlated axes between two groups of variables. Unit spiking data in *M1* and cerebellum from –2 s to + 2 s around task-start for each trial were binned at 50ms and concatenated across *early* and *late* trials separately. Our sessions contained at least 2 *M1* or cerebellar units. CCA models were then fit using the MATLAB function *canoncorr*. This function involves transforming the data to have zero mean and unit standard deviation prior to computing canonical variables (CVs). The number of CVs determined by the function is equal to the minimum number neurons in *M1* or cerebellum in a session.

To determine which CVs were significant, the canonical correlation of each CV was compared with a bootstrap distribution made of the canonical correlation of top CV from CCA models fit to trial-shuffled data (10^4^ shuffles). Specifically, before fitting CCA, trials from cerebellum were concatenated in the order in which they occurred, while trials from M1 were randomly permuted prior to concatenation. This method maintains local neural activity structure but breaks trial-by-trial relationship between neural modulation between these two regions. This provides a floor for the degree of correlation expected from the fact that many units in both regions have firing rate fluctuations around task-start. A CV was considered significant if its canonical correlation was greater than the 95^th^ percentile of the bootstrap distribution. All sessions had 1–2 significant CVs. For evaluating cross-area signals, only the top CV was used. We performed the CCA analysis with spiking activity from an equal number of successful trials from *early* and *late* learning.

#### Regression analysis

We use generalized linear model (GLM), using the MATLAB function *fitglm* to predict *M1* BMI potent space activity from cerebellum *TR_i_*’ s. For predicting *M1 TR_d_* activity from cerebellum *TR_i_*’s, BMI potent space activity was calculated as the difference between summed *M1 TR_d+_* activity & summed *M1 TR_d–_* activity (+/– being +ve or –ve unit weight associated *TR_d_* ’s), which was used as the response variable^44^. Predictors were binned firing rates of cerebellum *TR_i_* units, where each neuron appeared more than once with variable time lags ranging from –50ms to +50ms relative to the BMI–potent activity. Such horizontally stacked neural data corresponding to each trial in a session was used as predictive variable. In every session, for each model, a cross-validated *R*^2^ value was computed by splitting each session data into 9– folds for training and 1–fold for test which was repeated 10 times. *R*^2^ values were computed between the true response variable and the model output. The *R*^2^ values reported are the average across all 10 combinations of testing/training data. For predicting *M1 TR_i_* and *M1 TU* activity from cerebellum *TR_i_* ’s, a *‘surrogate BMI–potent space’* was created from *M1* neural activity by randomly selecting matched numbers of task–indirect/ unrelated units (*M1 TR_i_ / M1 TU*) to stand in for the true direct units (*M1 TR_d_ ’s*). The difference of summed activity in +ve and –ve pools obtained was used as the response variable. This process was repeated for 50 choices of such units per dataset.

#### Video tracking and analysis

We performed automated tracking of the tip of the feeding tube, the forepaws, and the head of the rats using DeepLabCut^79^. We performed cross-correlation between the trajectories of the feeding tube and either forepaw/ head using *corrcoef* function of MATLAB and looked at correlation coefficient *R* and *p* value for every trial (**Extended Fig. 2**).

#### Feeding tube trajectory analysis

We analyzed the feeding tube trajectory to look at its angular position and velocity from *early* to *late* trials. We looked at the angular position by plotting the x and y position of the feeding tube in the camera field of view over time, for *early* and *late* trials. We calculated the instantaneous angular velocity for every trial by looking at the displacement of the feeding tube between the subsequent frames and dividing it with the elapsed time. For every *early* and *late* trial, we calculated speed of the feeding tube by looking at the total displacement of the feeding tube from *P_1_* to *P_2_* and dividing it with the time to task completion for that trial. We then took the average speed for *early* and *late* trials. For speed consistency analysis, we interpolated feeding tube trajectory from *early* and *late* trials separately to make it equal to the feeding tube trajectory of the longest trial in each condition. We then calculated feeding tube velocity on these interpolated trials. We correlated the velocity profiles of individual *early* and *late* trials to the template velocity profile (constructed form the mean of velocity profiles of *late* trials) to show the change in speed consistency.

#### Statistical analysis

All statistical analyses were implemented within MATLAB. The linear mixed-effects model (implemented using MATLAB *fitlme*) was used to compare the differences in time to task completion, *MD_Δ_* (of different classes of cells from *early* to *late*), trial to mean correlation for speed consistency of the feeding tube, average speed of the feeding tube, M1/ cerebellum LFP power, *M1*–cerebellum LFP coherence, canonical correlations (*M1 TR*–cerebellum *TR_i_*, *M1 TR_d_*–cerebellum *TR_i_*, *M1 TR_i_*–cerebellum *TR_i_*, and *M1 TU*–cerebellum *TR_i_*) and on the *R*^2^ derived through spike GLM models, unless specified otherwise. This model accounts for the fact that units or sessions from the same animal are more correlated than those from different animals and is more stringent than computing statistical significance over all units and sessions^6, 7, 43, 69, 80^. We fitted random intercepts for each rat and reported the *p*-values for the regression coefficients associated with sessions, channels, or units. We also performed Kruskal-Wallis H-test with multiple comparisons in **Fig. 3e,f** and in **Extended Fig. 4**.To test the difference between two distributions, we did Kolmogorov–Smirnov two-sample test in **Fig. 4c,d,e,f**, **Extended Fig. 8a** and **Extended Fig. 9a**.

## Author Contributions

A.A. and T.G. designed the study. A.A., D.W.B., A.W.F. and N.P.D. carried out the electrophysiology experiments. A.A., D.W.B., A.W.F., N.P.D. and T.G. performed the surgical procedures. A.A. carried out all the analysis. R.R. made the predictive model and performed the regression analysis. D.W.B., A.W.F. and N.P.D. assisted with analysis. D.W.B., A.W.F. and N.P.D. carried out histology. A.A., R.R. and T.G. wrote the manuscript. D.W.B. edited the manuscript.

## Acknowledgment

The study was supported by American Heart Association (AHA; postdoctoral fellowship 897265 to A.A., predoctoral fellowship 1018175 to R.R., and career development award 847486 to T.G.), National Institutes of Health (NIH)’s National Institute for Neurological Disorders and Stroke (NINDS) (R00NS097620 and R01NS128469 to T.G.), National Science Foundation (award 2048231 to T.G.) and Cedars-Sinai Medical Center. A.A. also received support through Cedars-Sinai’s Center for Neural Science and Medicine postdoctoral fellowship.

---

**Extended Figure 1.**
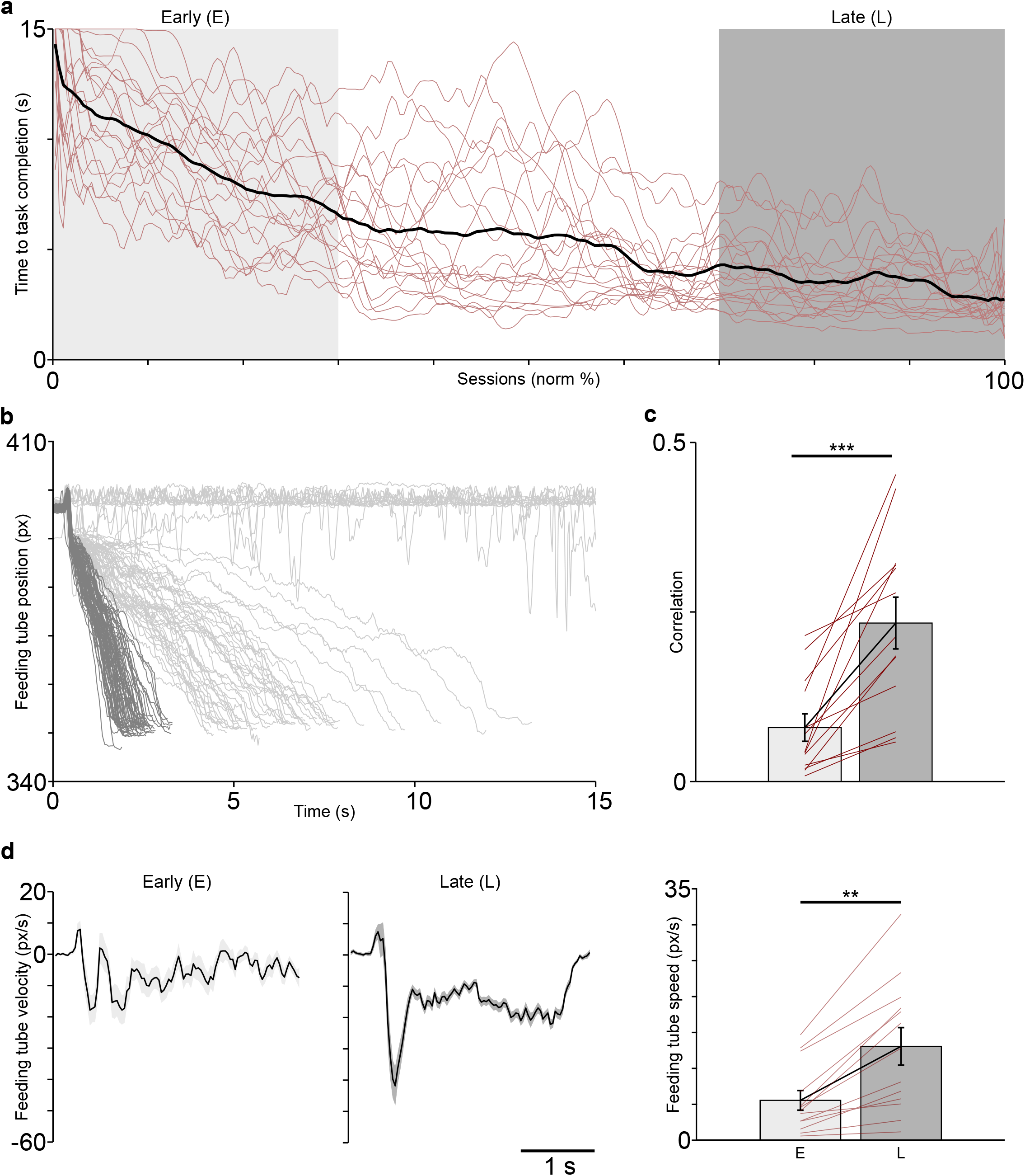
Learning curves and feeding tube trajectory. ***a,*** Learning curves from all the sessions depicted in red lines. Average learning curve is overlaid in black ***b,*** Position of the feeding tube from *early* (light gray) to *late* trials (dark gray) in representative session. ***c,*** Single-trial velocity trajectory correlation with the mean trajectory of *late* trials across sessions. ***d,*** Velocity of the feeding tube (*left*) and average speed of the feeding tube (*right*) from *early* to *late* trials. **p<0.01 ***p<0.001.

**Extended Figure 2.**
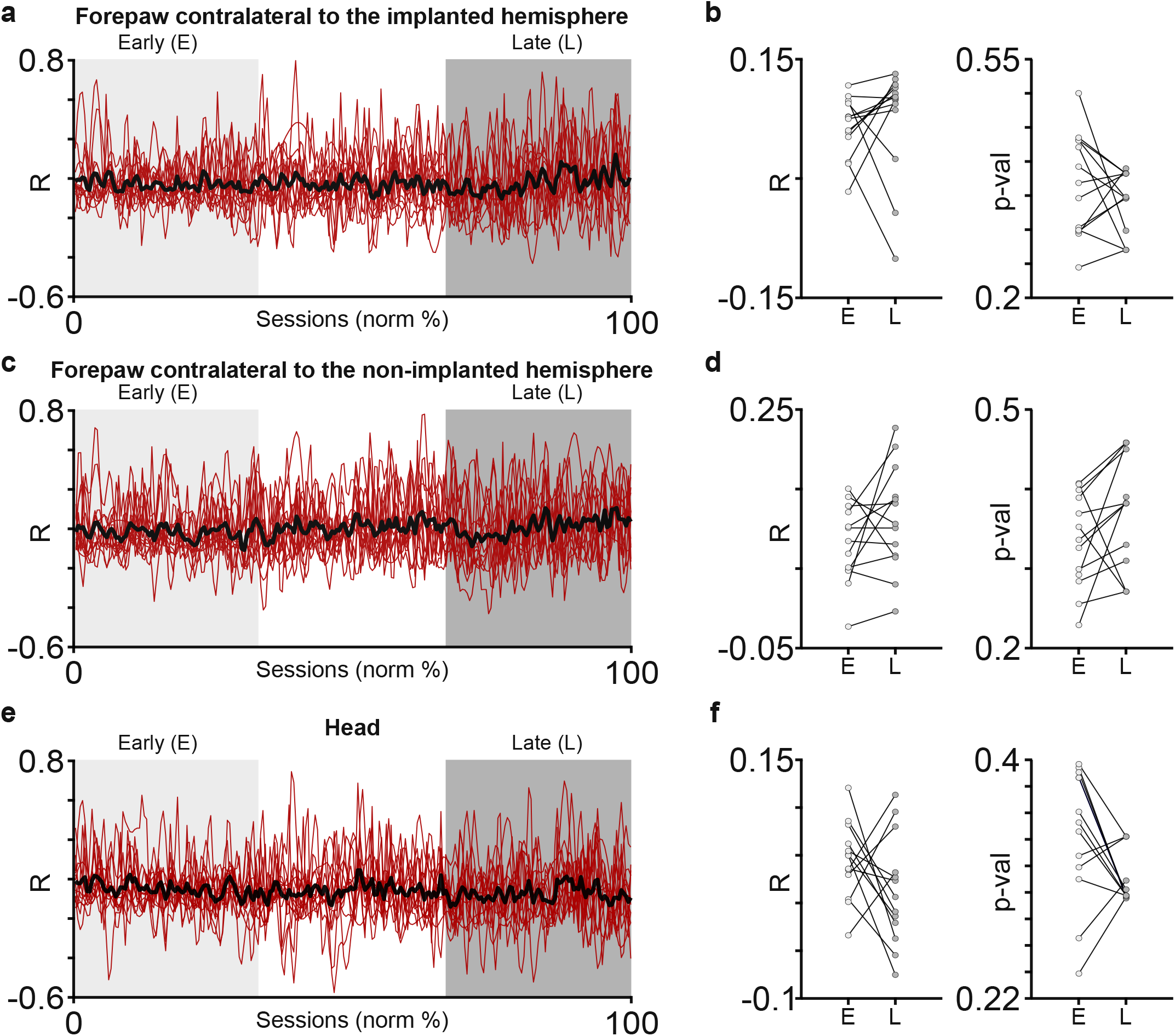
Movement correlation between body parts and feeding tube. ***a,*** Correlation between the forepaw contralateral to the implanted hemisphere and the feeding tube across trials during an example session. ***b,*** Correlation score (*left*) and p-value across (*right*) from *early* to *late* trials across sessions for forepaw contralateral to the implanted hemisphere indicating a non-significant trend for any correlated movement. ***c,*** Same as ***a*** for forepaw contralateral to the non-implanted hemisphere. ***d,*** Same as ***b*** for forepaw contralateral to the non-implanted hemisphere. ***e,*** Same as ***a*** for head. **f,** Same as ***b*** for head movements.

**Extended Figure 3.**
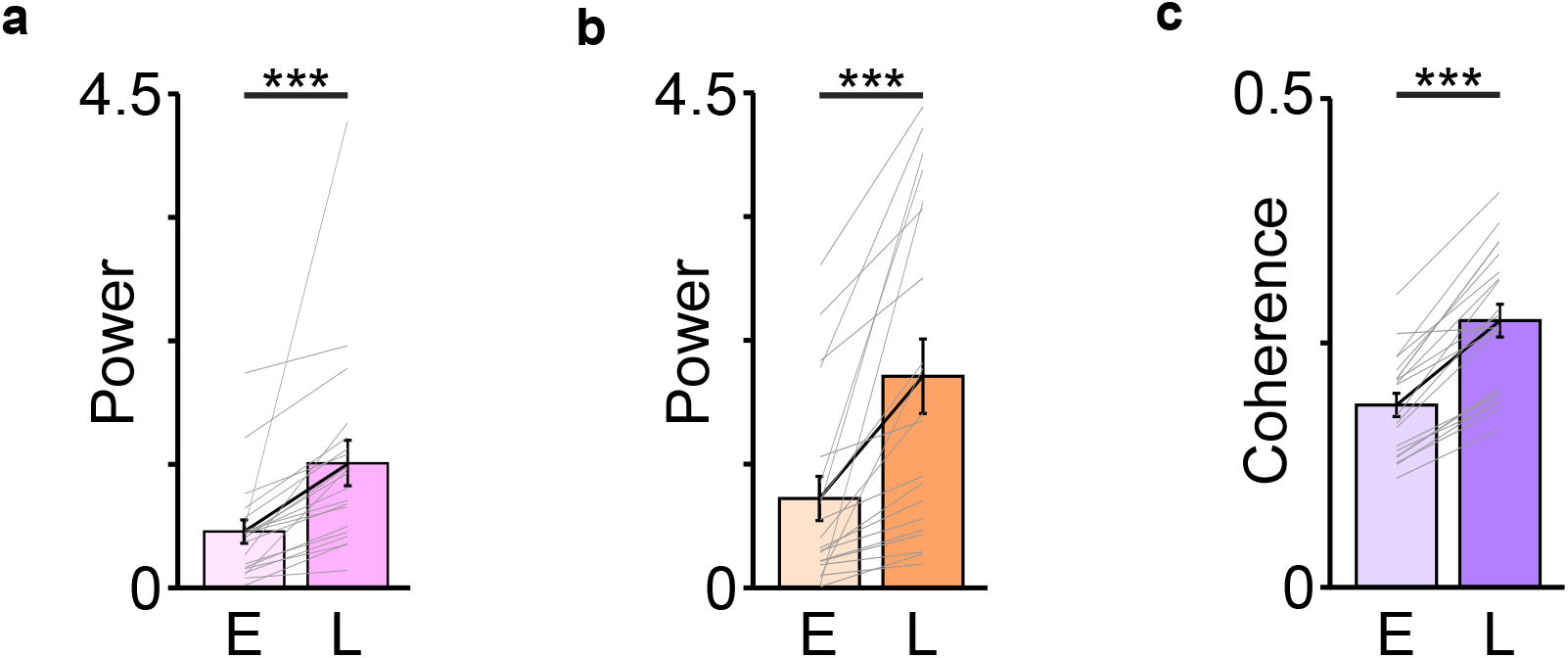
Co-ordinated task-related oscillations emerge in M1 and cerebellar LFPs across *early* (E) to *late* (L) successful only trials. ***a,*** Increase in the 3–6 Hz M1 LFP power across sessions. ***b***, Same as ***a*** for cerebellum LFP power. ***c***, Change in 3–6 Hz coherence from *early* to *late* successful trials across sessions. ***p<0.001.

**Extended Figure 4.**
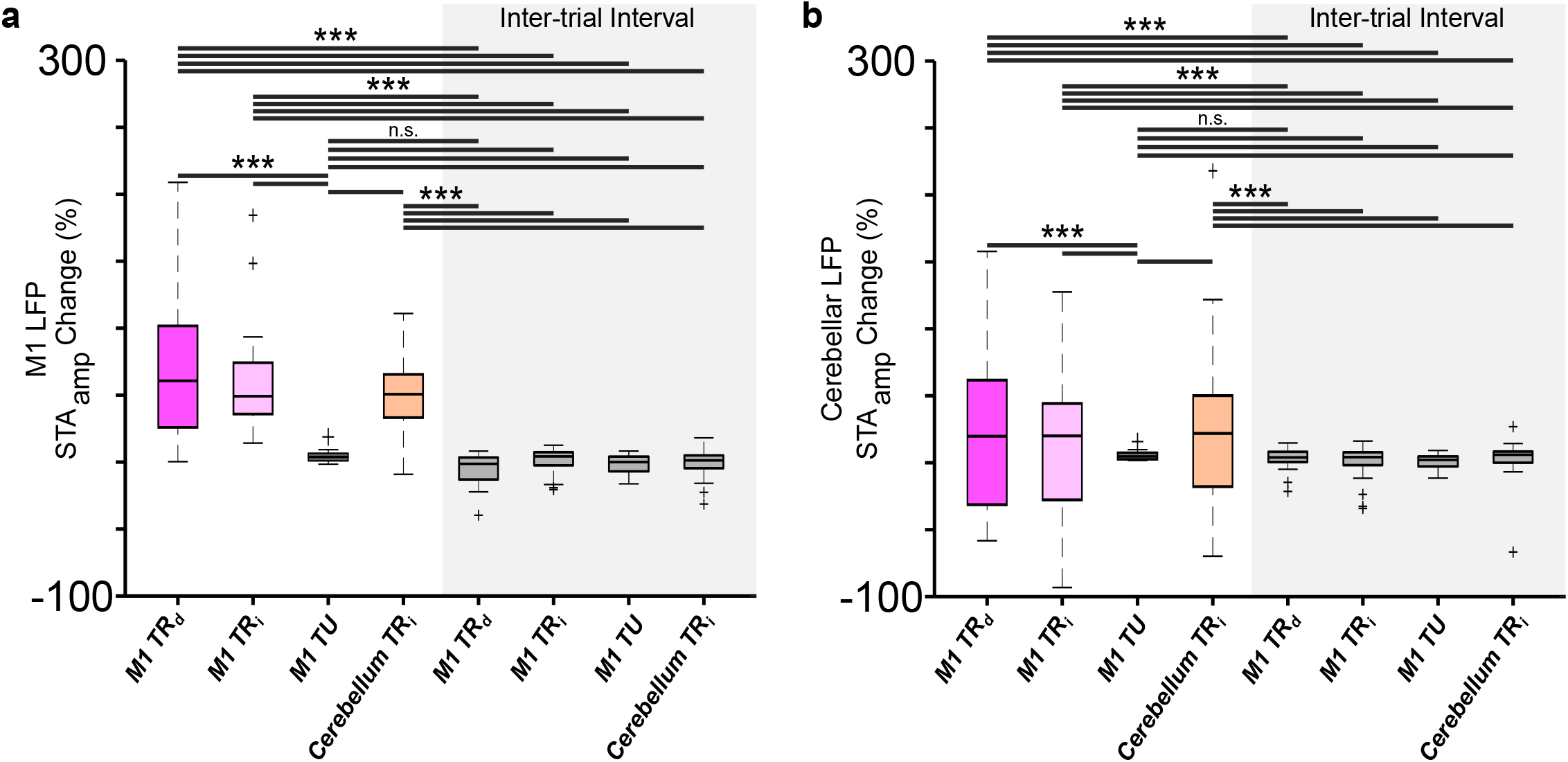
*M1* and cerebellum spike–LFP locking in only successful trials. ***a,*** Box–plot of percentage change in STA amplitude for *M1* LFP in each of the categories of units (box-plot conventions are same as **Fig. 3c**). ***b,*** Same as ***a*** for STA of cerebellum LFP. ***p<0.001. n.s.: non-significant.

**Extended Figure 5.**
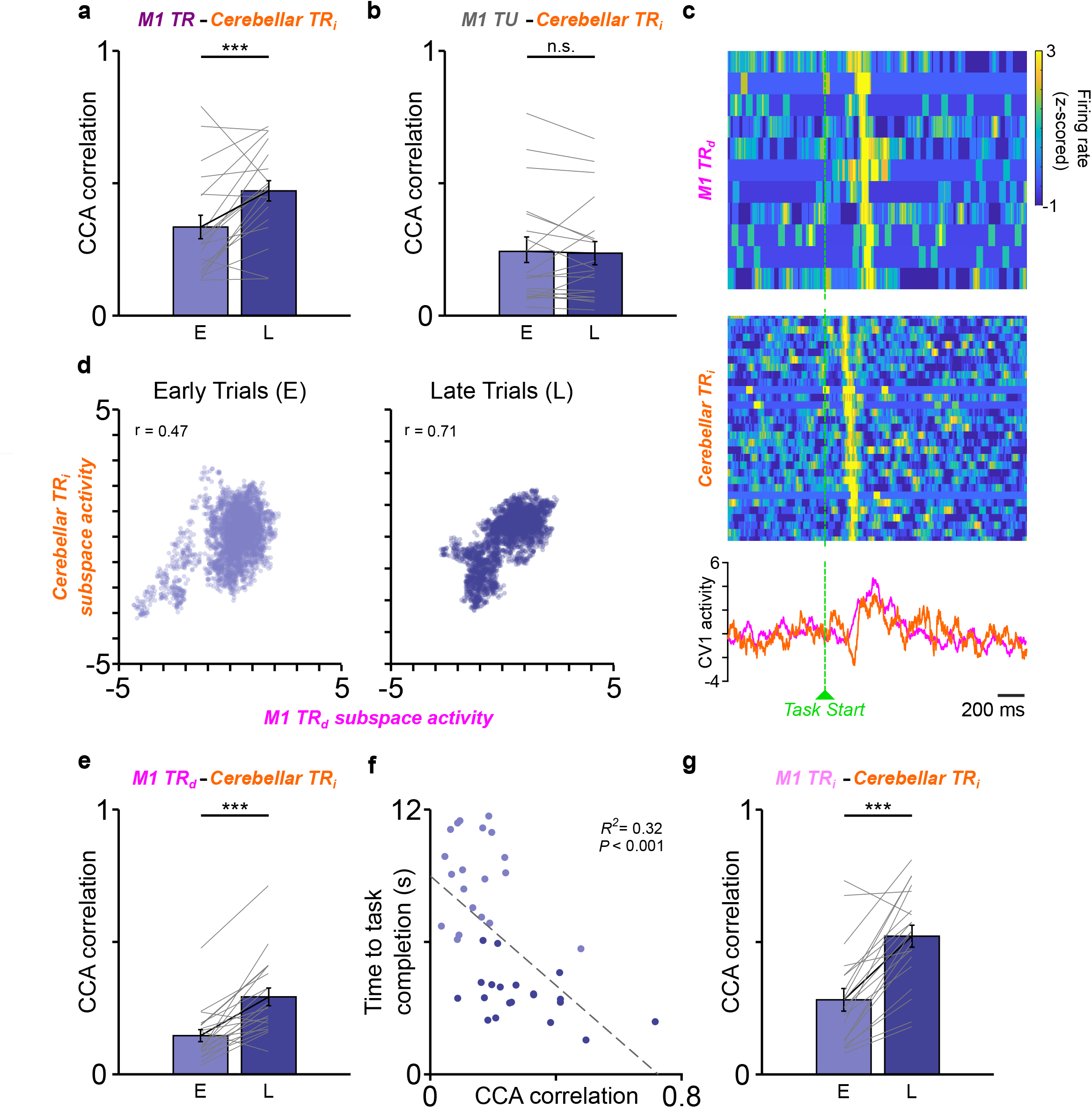
CCA correlation. ***a,*** Change in the canonical correlation score from only successful *early* to *late* trials across all sessions for *M1 TR* and cerebellar *TR_i_* units. ***b,*** Same as ***a*** for *M1 TU* and cerebellar *TR_i_* units. ***c,*** Single-trial *M1 TR_d_* and cerebellar *TR_i_* spiking activity along with the single-trial activations along CV1 around task-start. ***d,*** *M1 TR_d_* and cerebellum *TR_i_* subspace activity (from the CV1) around task-start (−2 to 2 s) for an example session. Each dot represents one time bin of *early* or *late* trials from the session. Canonical correlation score is given by *r*. ***e,*** Change in the canonical correlation score from *early* to *late* trials across all sessions for *M1 TR_d_* and cerebellar *TR_i_* units. ***f,*** Relationship between CCA correlation score of *TR_d_* and cerebellar *TR_i_* and average time to a successful trial across all sessions. ***g,*** Same as ***e*** for *M1 TR_i_ and* cerebellar *TR_i_* units. (***: p<0.001; n.s.: non-significant, p>0.05).

**Extended Figure 6:**
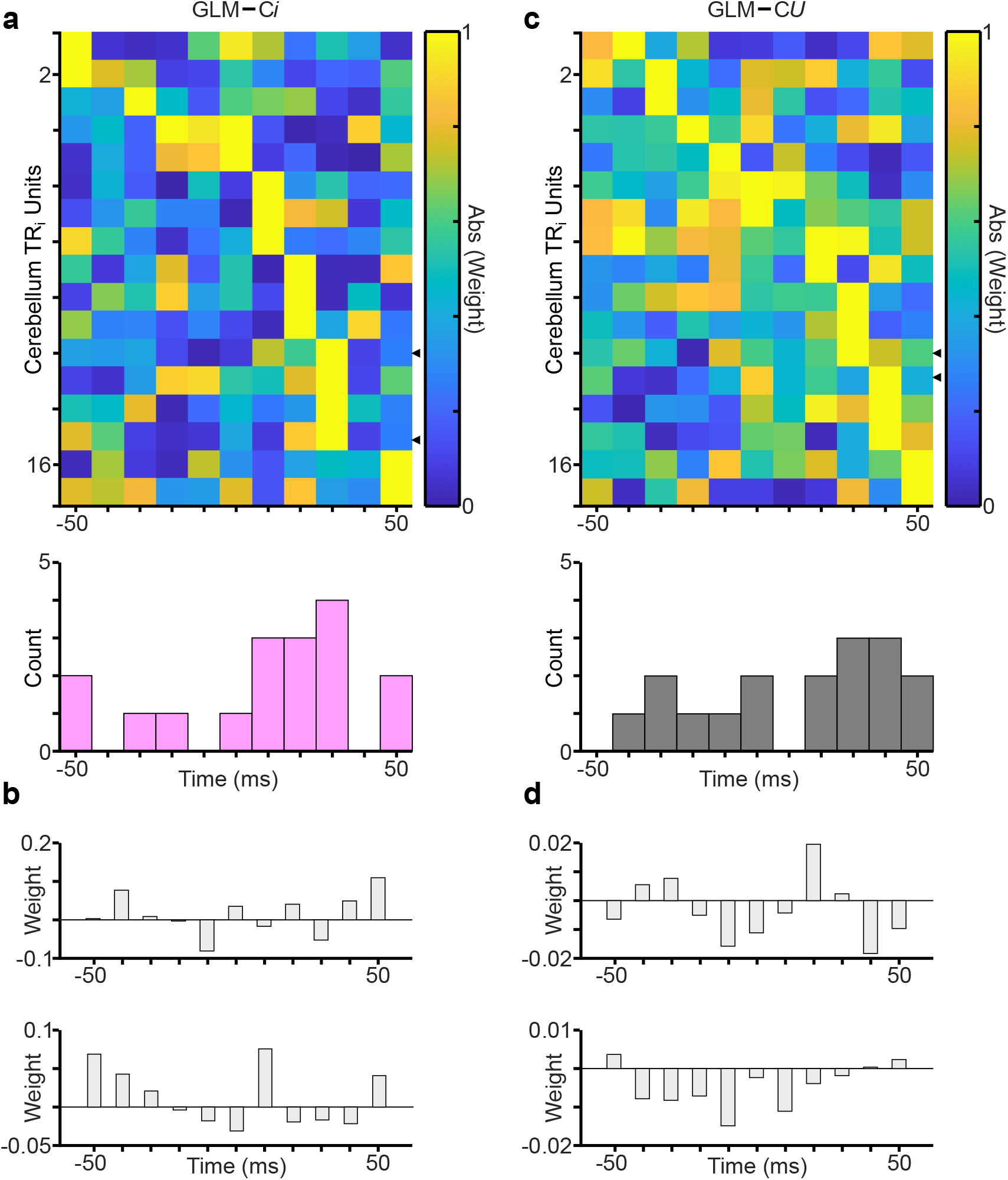
Timescale of M1 activity prediction from cerebellum *TR_i_* neural activity. ***a,*** Distribution of regression weight magnitude in one example session for GLM–C*i* (fitted to neural data binned at 10ms). *Top*, for each cerebellum *TR_i_* unit, regression weights were assigned for a variety of time lags and the absolute value of those weights is indicated by color. Units are sorted according to the time of the largest magnitude weight. Tick marks on the right edge indicate the units shown in ***b***. *Bottom*, Histogram of the *τ* values with the largest magnitude weight for this dataset. ***b***, Example non-normalized weights for two cerebellum *TR_i_* neurons from one example session (neural data binned at 10ms). Height of bars indicates weights for example neurons at different time lags (*τ*) relative to the *M1* BMI-potent activity, with negative *τ* values meaning that cerebellum leads. ***c*** & ***d***, same as ***a*** & ***b*** but for GLM–C*U* model.

**Extended Figure 7.**
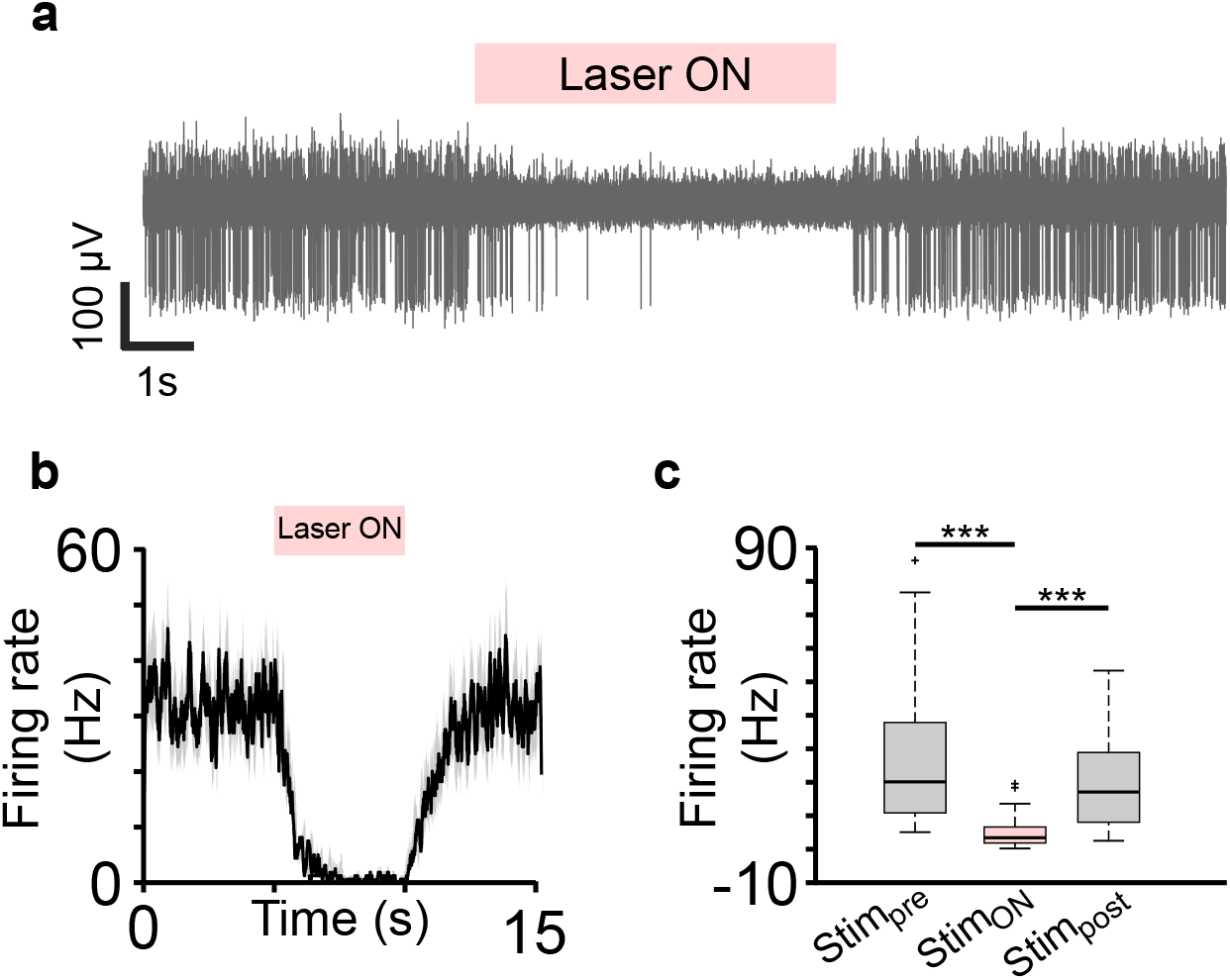
Inhibition of cerebellar cortical neural activity with optogenetic illumination. ***a,*** A raw trace from an example channel showing reduction in cerebellar activity when the laser was turned on. ***b,*** PETH of an example unit showing reduction in firing rate with Laser ON. ***c,*** Box–plot showing change in the firing rate of all the recorded units *before* (*Stim_pre_*), *during* (*Stim_ON_*) and *after* (*Stim_post_*) cerebellar inhibition (box–plot conventions are same as **Fig. 3c**). ***p<0.001.

**Extended Figure 8.**
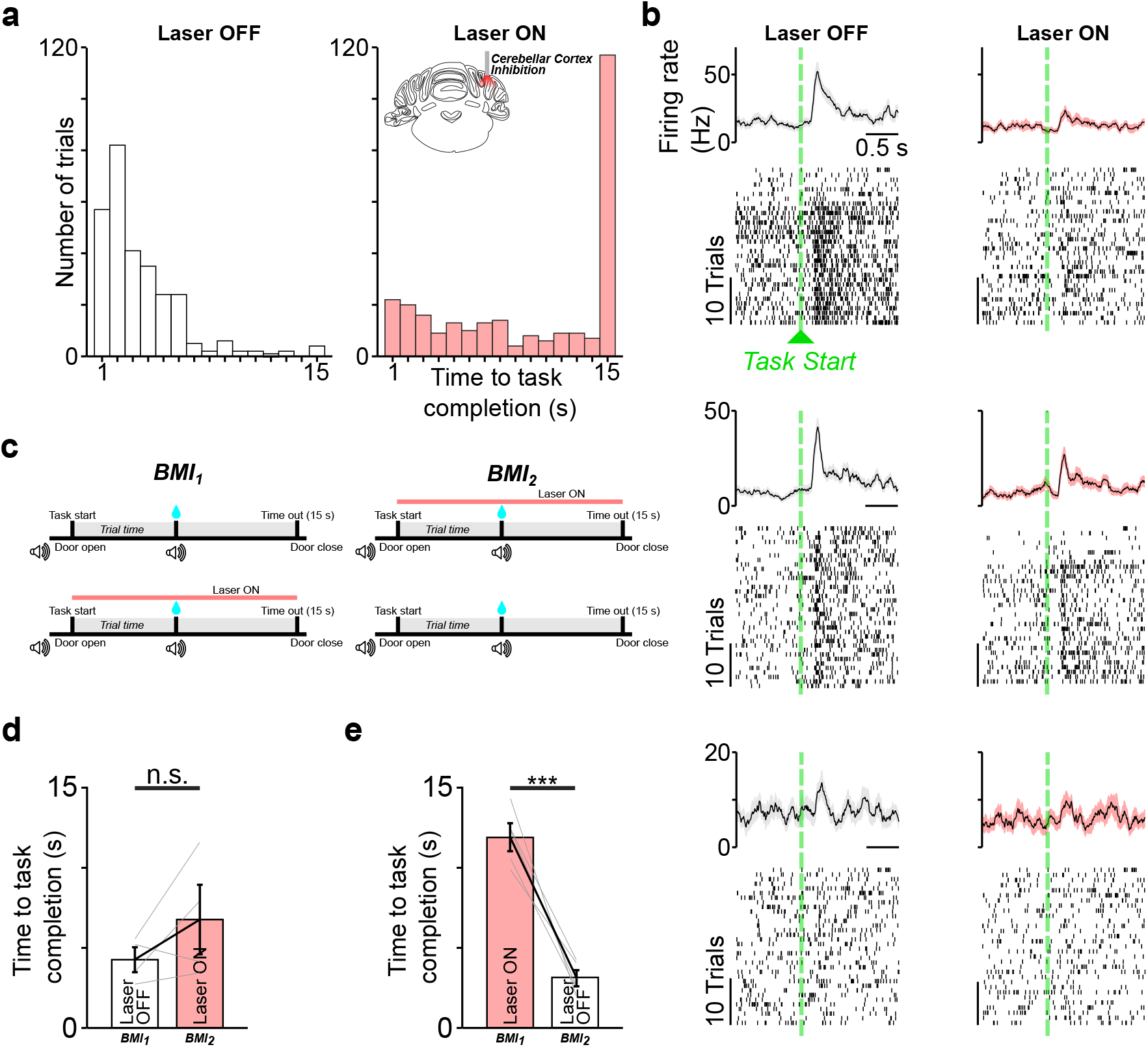
Effect of cerebellar cortex inhibition on neuroprosthetic control and M1 *‘direct’* activity. ***a,*** Distribution of time to task completion during *late* trials in Laser OFF (*left*) and Laser ON (*right*) condition. ***b,*** Example PETH and raster plots of three *M1 TR_d_* units during *late* trials in Laser OFF (*left*) and Laser ON (*right*) condition. ***c,*** Schematic of session structure. On a day (*top*), initial session was performed without cerebellar cortex inhibition trials (Laser OFF) followed by a subsequent session with cerebellar cortex inhibition (Laser ON). Next day (*bottom*), the order was inverted with initial session with cerebellar cortex inhibition performed first followed by a subsequent session without cerebellar cortex inhibition. ***d*** and ***e***, An increase in the time to task completion was observed irrespective of the order of cerebellar cortex stimulation, but it was significantly increased when inhibition occurred in *BMI_1_*. ***p<0.001; n.s.: not significant p>0.05.

**Extended Figure 9.**
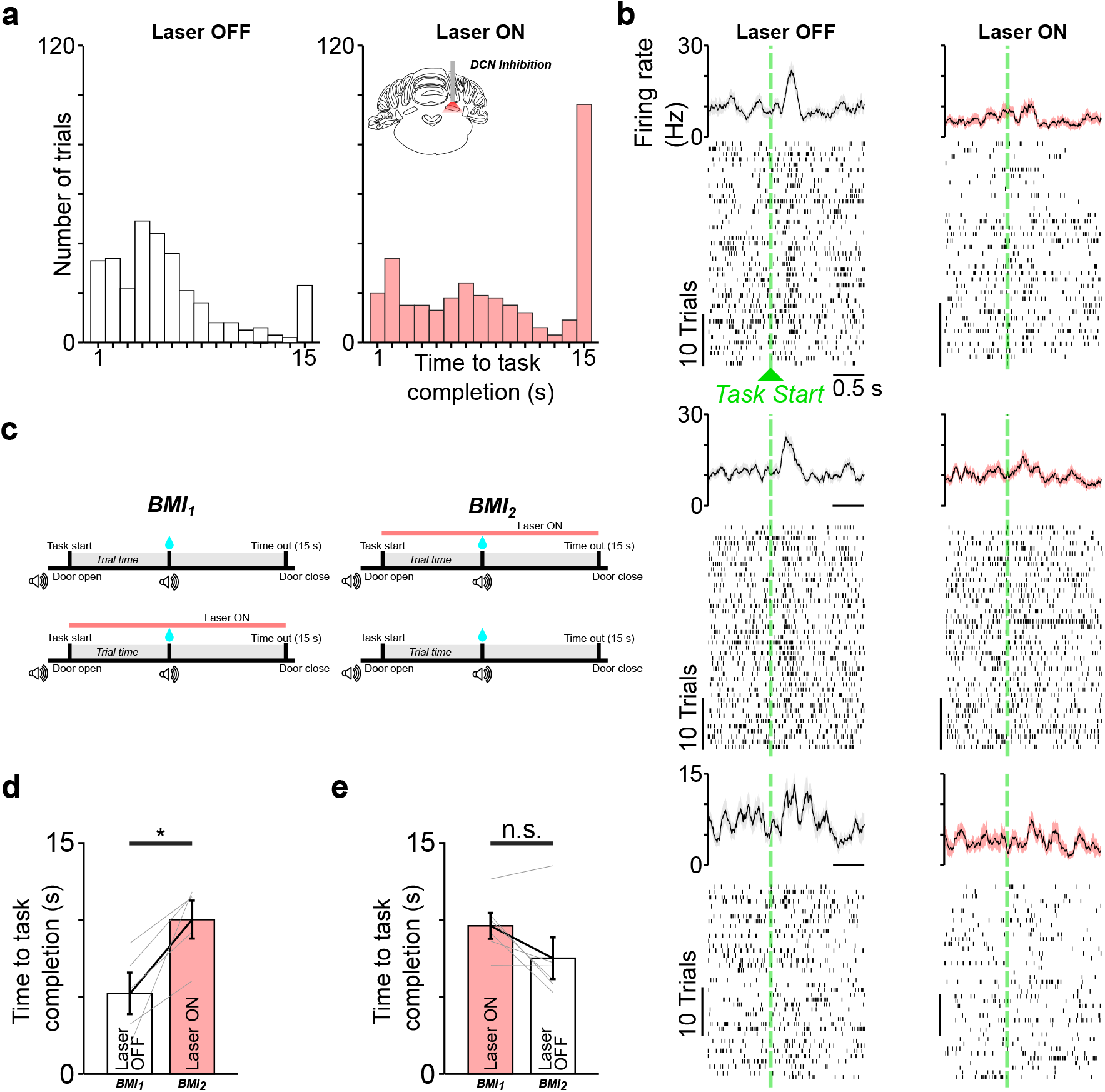
Effect of *DCN* inhibition on neuroprosthetic control and M1 *‘direct’* activity. ***a,*** Distribution of time to task completion during *late* trials in Laser OFF (*left*) and Laser ON (*right*) condition. ***b,*** Example PETH and raster plots of three *M1 TR_d_* units during *late* trials in Laser OFF (*left*) and Laser ON (*right*) condition. ***c,*** Schematic of session structure. On a day (*top*), initial session was performed without *DCN* inhibition trials (Laser OFF) followed by a subsequent session with *DCN* inhibition (Laser ON). Next day (*bottom*), the order was inverted with initial session with *DCN* inhibition performed first followed by a subsequent session without *DCN* inhibition. ***d*** and ***e***, An increase in the time to task completion was observed irrespective of the order of *DCN* stimulation, but it was significantly increased when inhibition occurred in *BMI_2_*. *p<0.05; n.s.: not significant p>0.05.

**Extended Figure 10.**
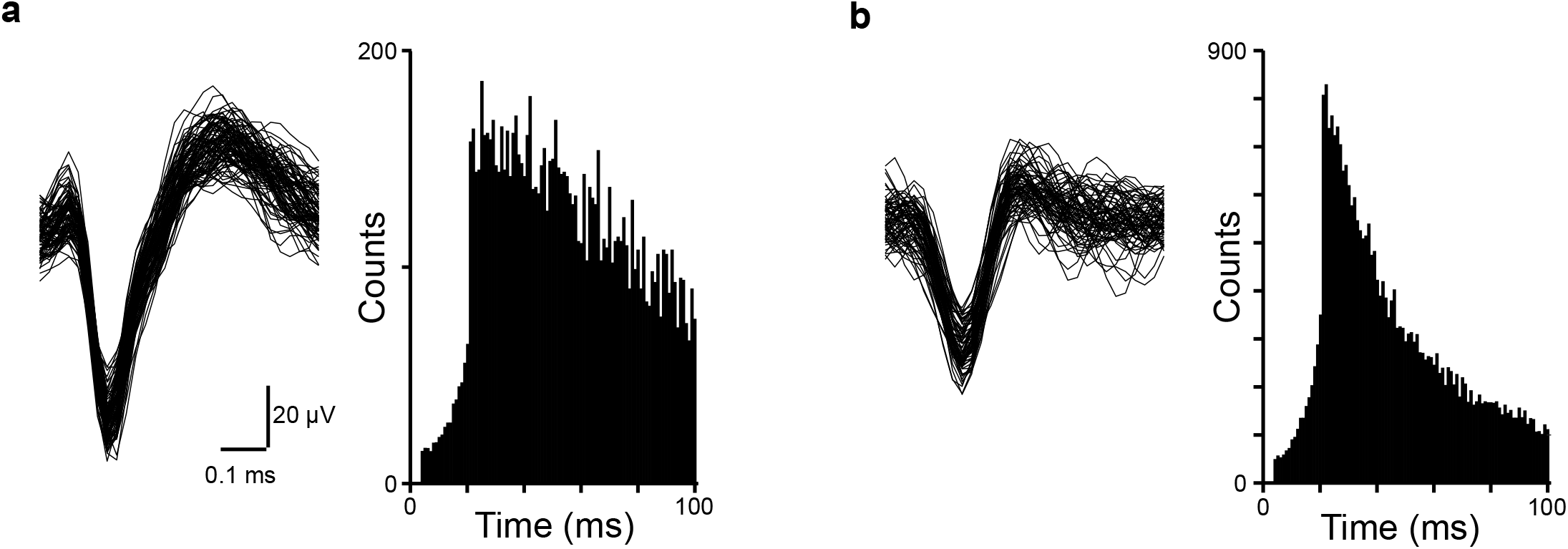
Single unit waveforms and inter-spike interval (ISI). ***a,*** Left panel shows waveforms of a representative *M1 TR_d_* unit. Right panel shows ISI histograms for the same unit. *b,* Left panel shows waveforms of a representative task–related *M1 TR_i_* unit. Right panel shows ISI histograms for the corresponding units in the left panel.

**Extended Figure 11.**
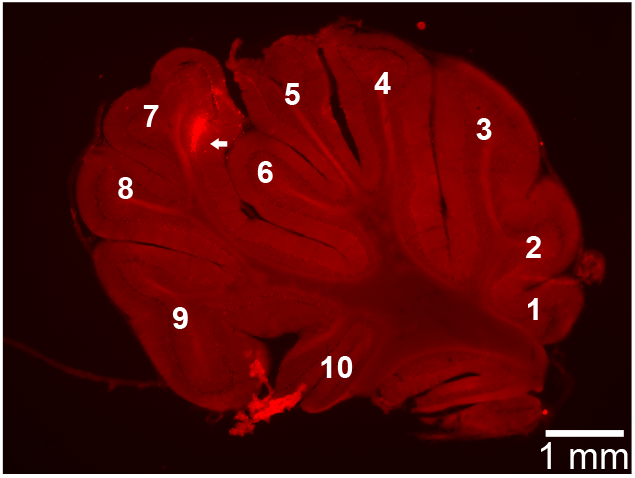
Cerebellum electrode localization. The location of the electrode in the cerebellum is indicated by a white arrow in the image of a brain section 4.75 mm lateral to the midline. Numbers indicate cerebellar lobule.

**Supplementary Table 1.**
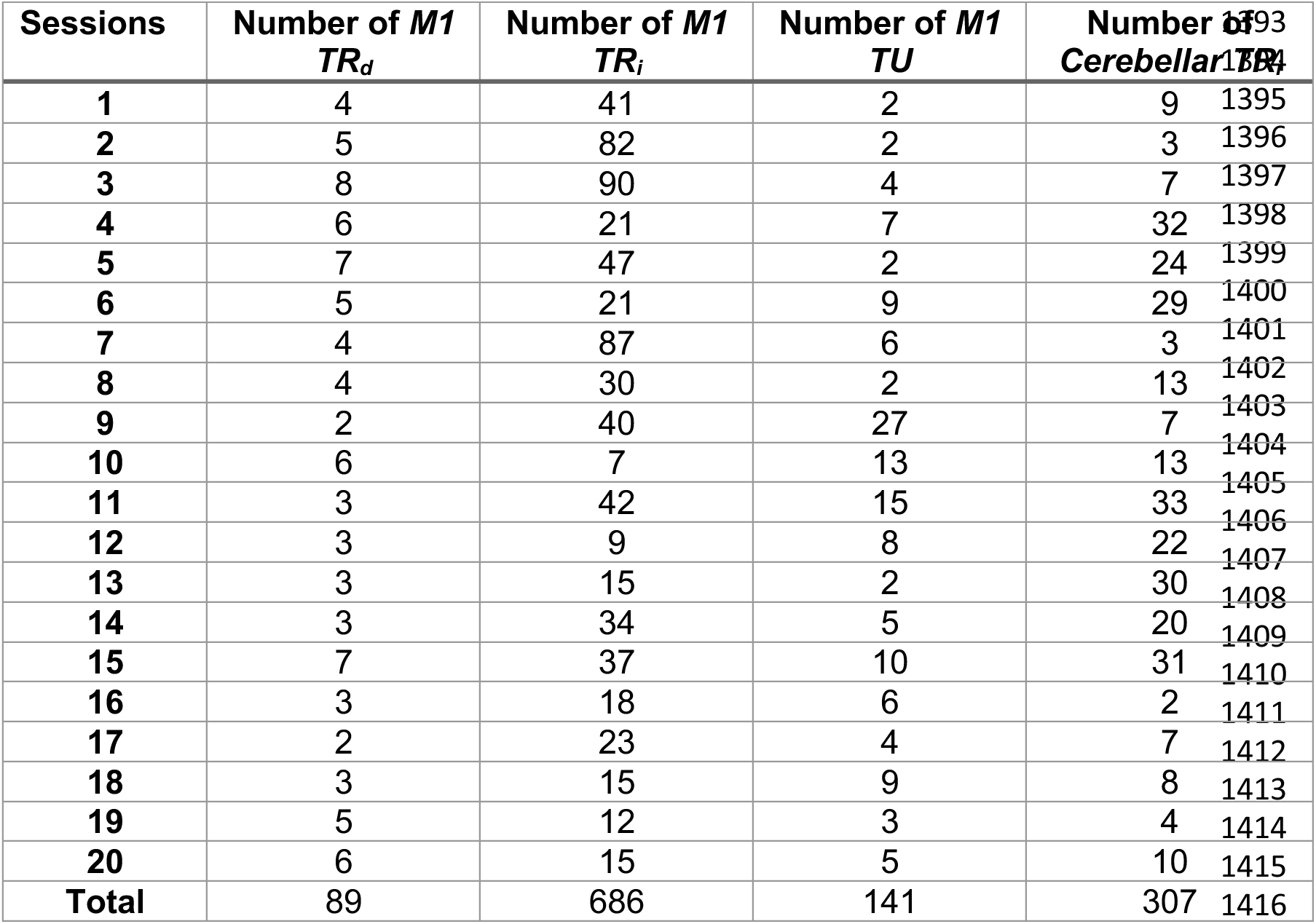
Number of *M1* and cerebellar units. Number of *M1* and cerebellar units of different classes across all neuroprosthetic sessions.

