## Supplementary Table 1 for "Cortico-cerebellar coordination facilitates neuroprosthetic control"

| Sessions | Number of *M1 TR_d_* | Number of *M1 TR_i_* | Number of *M1 TU* | Number of *Cerebellar TR_i_* |
| --- | --- | --- | --- | --- |
| 1 | 4 | 41 | 2 | 9 |
| 2 | 5 | 82 | 2 | 3 |
| 3 | 8 | 90 | 4 | 7 |
| 4 | 6 | 21 | 7 | 32 |
| 5 | 7 | 47 | 2 | 24 |
| 6 | 5 | 21 | 9 | 29 |
| 7 | 4 | 87 | 6 | 3 |
| 8 | 4 | 30 | 2 | 13 |
| 9 | 2 | 40 | 27 | 7 |
| 10 | 6 | 7 | 13 | 13 |
| 11 | 3 | 42 | 15 | 33 |
| 12 | 3 | 9 | 8 | 22 |
| 13 | 3 | 15 | 2 | 30 |
| 14 | 3 | 34 | 5 | 20 |
| 15 | 7 | 37 | 10 | 31 |
| 16 | 3 | 18 | 6 | 2 |
| 17 | 2 | 23 | 4 | 7 |
| 18 | 3 | 15 | 9 | 8 |
| 19 | 5 | 12 | 3 | 4 |
| 20 | 6 | 15 | 5 | 10 |
| Total | 89 | 686 | 141 | 307 |

**Supplementary Table 1. Number of *M1* and cerebellar units.** Number of *M1* and cerebellar units of different classes across all neuroprosthetic sessions.
